# Missing the sweet spot: one of the two N-glycans on human Gb3/CD77 synthase is expendable

**DOI:** 10.1101/2021.02.04.429793

**Authors:** Krzysztof Mikolajczyk, Anna Bereznicka, Katarzyna Szymczak-Kulus, Katarzyna Haczkiewicz-Lesniak, Bozena Szulc, Mariusz Olczak, Joanna Rossowska, Edyta Majorczyk, Katarzyna Kapczynska, Nicolai Bovin, Marta Lisowska, Radoslaw Kaczmarek, Arkadiusz Miazek, Marcin Czerwinski

## Abstract

N-glycosylation is a ubiquitous posttranslational modification that may influence folding, subcellular localization, secretion, solubility and oligomerization of proteins. In this study, we examined the effects of N-glycans on the activity of human Gb3/CD77 synthase, which catalyzes the synthesis of glycosphingolipids with terminal Galα1→4Gal (Gb3 and the P1 antigen) and Galα1→4GalNAc disaccharides (the NOR antigen). The human Gb3/CD77 synthase contains two occupied N-glycosylation sites at positions N_121_ and N_203_. Intriguingly, we found that while the N-glycan at N_203_ is essential for activity and correct subcellular localization, the N-glycan at N_121_ is dispensable and its absence did not reduce, but, surprisingly, even increased the activity of the enzyme. The fully N-glycosylated human Gb3/CD77 synthase and its glycoform missing the N_121_ glycan correctly localized in the Golgi, whereas a glycoform without the N_203_ site partially mislocalized in the endoplasmic reticulum. A double mutein missing both N-glycans was inactive and accumulated in the endoplasmic reticulum. Our results suggest that the decreased specific activity of human Gb3/CD77 synthase glycovariants results from their improper subcellular localization and, to a smaller degree, a decrease in enzyme solubility. Taken together, our findings show that the two N-glycans of human Gb3/CD77 synthase have opposing effects on its properties, revealing a dual nature of N-glycosylation and potentially a novel regulatory mechanism controlling the biological activity of proteins.

## Introduction

Glycosyltransferases (GTs) constitute a large group of enzymes catalyzing transfer of sugar residues from carbohydrate donors (*e.g.* UDP-Gal, CMP-NeuNAc) to diverse acceptor molecules, forming glycosidic bonds with retention or inversion of the attached sugar configuration [Breton C., Fournel-Gigleux S. et al. 2012]. On the basis of structural analysis most GTs are classified into five overall folds: GT-A [Taujale R., Venkat A. et al. 2020], GT-B [Albesa-Jové D., Giganti D. et al. 2014], GT-C [Bohl T., Bai L. et al 2021], GT-D [Zhang H., Zhou M. et al. 2016] and GT-E [Kattke M.D., Gosschalk J.E. et al. 2019].

N-glycosylation is a ubiquitous posttranslational modification (PTM) of proteins; an estimated 50% may be N-glycosylated [Goettig P., 2016]. N-glycans are attached to a protein by asparagine within a canonical N-X-S/T motif called a sequon (where X is any amino acid residue except proline); however, other sequons, such as N-X-C, N-Q-C, N-S-G or Q-G-T sequons (referred to as noncanonical) may also be utilized [Lowenthal M.S., Davis K.S. et al. 2016]. In eukaryotes, biosynthesis of N-glycans occurs in two cellular compartments: (1) the endoplasmic reticulum (ER), in which the dolichol-P-linked oligosaccharide substrate is attached to the asparagine, and (2) the Golgi apparatus, which contains glycan-processing enzymes involved in trimming and maturation of N-glycans [Stanley P., Taniguchi N. et al. 2017]. Glycosylation may play a major role in many cellular processes, such as protein folding [Jayaprakash N.G., Surolia A. et al. 2017], maturation [Agthe M., Garbers Y. et al. 2018], secretion [Fiedler K., Simons K. et al. 1995], intracellular trafficking [Rosnoblet C., Peanne R. et al. 2013], cell-cell interactions [Varki A., 2017], immune responses [Ryan S.O., Cobb B.A. et al. 2012] and disease progression [Kizuka Y., Kitazume S. et al. 2017; B.N., Patel P.S. et al. 2017].

Human Gb3/CD77 synthase (α1,4-galactosyltransferase, P1/P^k^ synthase, EC 2.4.1.228) encoded by the *A4GALT* gene, is a type II transmembrane GT with C-terminal globular catalytic domain facing the Golgi lumen and N-terminal cytoplasmic domain; it belongs to the CAZy glycosyltransferase family 32 (Carbohydrate Active Enzymes database, CAZy, http://www.cazy.org/) [Lombard V., Golaconda Ramulu H. et al 2014]. The enzyme retains the donor’s anomeric carbon stereochemistry after glycosidic bond formation [Furukawa K., Kondo Y. et al. 2014; Okuda T., Tokuda N. et al. 2006]. *In silico* analysis predicted that human Gb3/CD77 synthase adopts the GT-A fold structure (CAZy, http://www.cazy.org/). Members of the GT-A superfamily typically require a divalent metal ion (usually Mn^2+^) at the catalytic center, which is coordinated by two aspartic acid residues, creating a DXD motif (D_192_TD in human Gb3/CD77 synthase according to UniProt Q9NPC4, https://www.uniprot.org/).

Previously, we found that in addition to the high-frequency *A4GALT* gene variant (GenBank NG_007495.2), there is another variant with a point mutation c.631C>G (rs397514502, GenBank NG_007495.2) giving rise to the protein with p.Q211E substitution (hereafter referred to as E or mutein). The enzyme encoded by the high-frequency gene (here designated as Q) catalyzes the transfer of galactose from UDP-galactose to lactosylceramide or paragloboside, producing globotriaosylceramide (Gb3, P^k^, CD77) and the P1 antigen, respectively. Both Gb3 and P1 terminate with a Galα1→4Gal moiety. It was recently shown that Gb3/CD77 synthase can also add galactose to Galβ1→Cer, creating galabiosylceramide (Galα1→4Galβ1-Cer) [Akiyama H., Ide M. et al. 2021]. In addition, the mutein can transfer galactose to a terminal GalNAc, giving rise to the rare NOR antigen (carried by NOR1, NOR_int_ and NOR2 glycosphingolipids), which terminates with Galα1→4GalNAc structures [Suchanowska A., Kaczmarek R. et al. 2012]. The ability of human Gb3/CD77 synthase to use two different acceptors is a unique case of glycosyltransferase promiscuity [Kaczmarek R., Duk M. et al. 2016]. All these antigens (Gb3, P1 and NOR) belong to the human P1PK blood group system: the presence or absence of P1 on red blood cells (RBCs) determines the P_1_ or P_2_ blood group phenotypes, respectively. In the rare p phenotype, which may be caused by null mutations in *A4GALT*, the P1PK antigens are not detected on RBCs. The presence of NOR antigens results in the rare NOR phenotype [Kaczmarek R., Szymczak-Kulus K. et al. 2018; Kaczmarek R., Buczkowska A. et al. 2014].

Gb3 is present in human RBCs and lymphocytes, heart, lung, kidney, smooth muscle and epithelium of gastrointestinal tract [Cooling L., 2015]. Several reports showed that elevated expression of Gb3 was found in colorectal, gastric and ovarian cancers [Kovbasnjuk O., Mourtazina R. et al. 2005; Geyer P.E., Maak M. et al. 2016]. In Fabry disease, which is an X-linked lysosomal storage disorder caused by deficiency of α-galactosidase (OMIM 301500), Gb3 accumulates in organs throughout the body [Miller J.J., Kanack A.J. et al. 2020]. Expression of the P1 antigen seems to be limited to the erythroid lineage [Cooling L., 2015], but it was also detected on ovarian cancer cells, where it was designated as cancer-associated antigen [Jacob F., Anugraham M. et al. 2014].

Galα1→4Gal disaccharide-containing glycosphingolipids (GSLs), such as Gb3 are targeted by bacterial adhesins, including PapG of uropathogenic *Escherichia coli*, and viruses [Cooling L., 2015]. Gb3 is also the main receptor for Shiga toxins (Stxs) secreted by Shiga toxin-producing *E. coli* (STEC) [Lee M.S., Tesh V., 2019]. Humans contract STEC infections by ingestion of contaminated food or water; every year STEC cause an estimated 2.8 million severe illnesses worldwide [Majowicz S.E., Scallan E. et al. 2014]. Shiga toxins can cause hemorrhagic colitis, which may progress to hemolytic-uremic syndrome (HUS), an acute and often fatal complication [Cody E.M., Dixon B.P., 2019; Bruyand M., Mariani-Kurkdjian P. et al. 2018].

Several studies showed that elimination of N-glycans from GTs may affect their activity [Mikolajczyk K., Kaczmarek R. et al. 2020]. The protein sequence of human Gb3/CD77 synthase includes two potential N-glycosylation sites at positions N_121_ and N_203_ (UniProt Q9NPC4, https://www.uniprot.org/). Using site-directed mutagenesis, we generated six N-glycosylation variants of the human Gb3/CD77 synthase (three for Gb3/CD77 synthase and three for the mutein) with substituted N-glycosylation sequons, and analyzed their activity, subcellular trafficking and secretion in CHO-Lec2 cells transfected with vectors encoding the glycovariants. Finally, we evaluated the sensitivities of CHO-Lec2 cells expressing different glycovariants to Shiga toxins.

## Results

### Gb3/CD77 synthase contains two occupied N-glycosylation sites

Human Gb3/CD77 synthase contains two potential N-glycosylation sites: N_121_-A-S and N_203_-L-T (UniProt Q9NPC4, https://www.uniprot.org/) (Fig. 1). Our preliminary studies suggested that treatment of the Q enzyme with peptide-N-glycosidase F (PNGase F) abolished its *in vitro* activity (Fig. S1). Six glycovariants were generated: four single-mutants with p.S123A substitution (Q_S123A_ and E_S123A_ for Gb3/CD77 synthase and its mutein, respectively) and p.T205A substitution (Q_T205A_ and E_T205A_ for Gb3/CD77 synthase and its mutein, respectively), as well as two double-mutants (Q_S123A_/Q_T205A_ and E_S123A_/E_T205A_) (Fig. 1). The enzymes without any substitutions at N-glycosylation sites (fully N-glycosylated) are referred to as “Q_full_” for the Gb3/CD77 synthase and “E_full_” for the mutein.

**Fig. 1.**
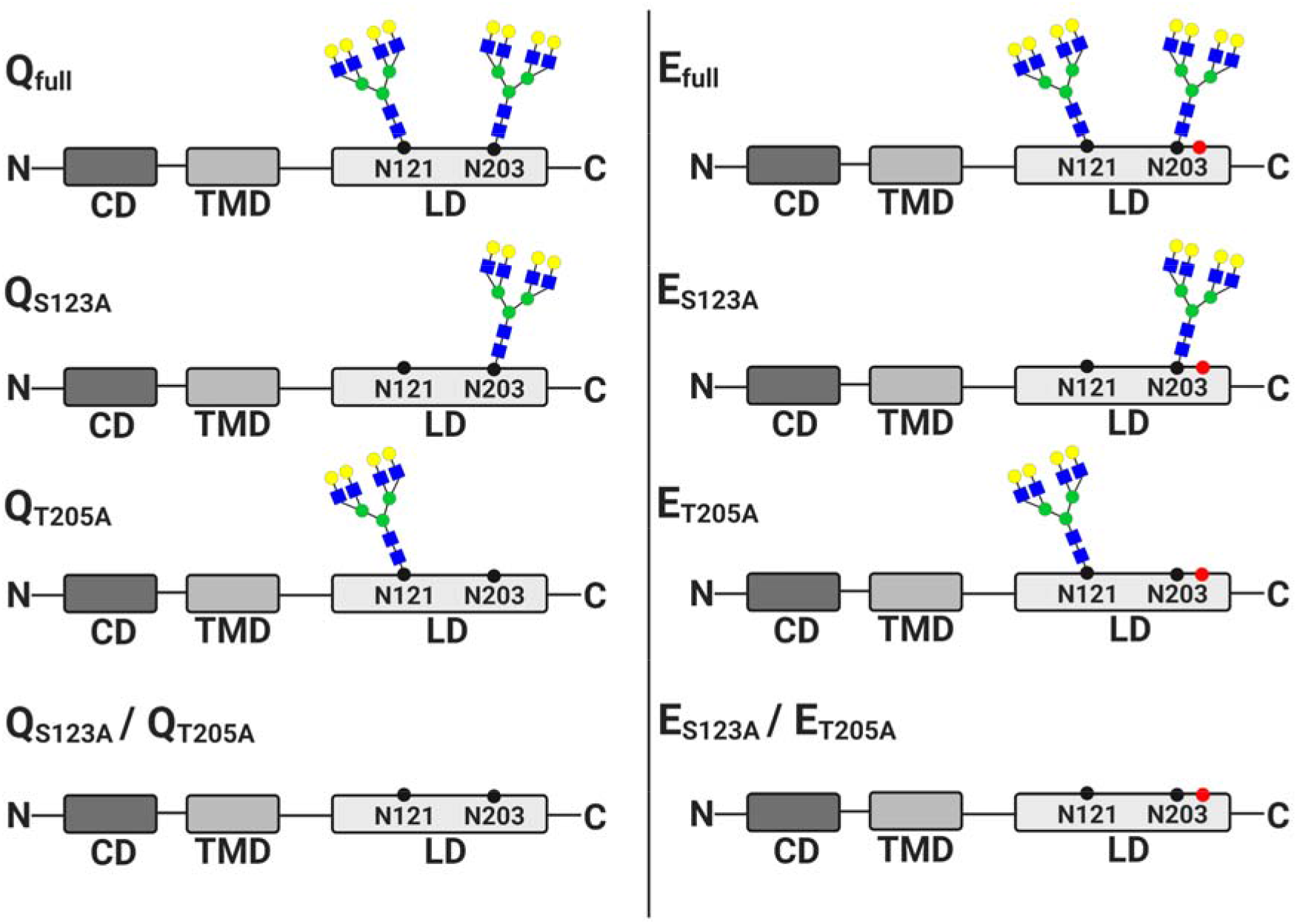
Schematic representation of human Gb3/CD77 synthase and its mutein glycovariants with N-glycosylation sites occupation. The enzyme contains cytoplasmic domain (CD, encompasses 1-22 amino acids residues), transmembrane domain (TMD, encompasses 23-43 amino acids residues) which resides the enzyme in Golgi apparatus membrane and lumenal domain (LD, encompasses 44-353 amino acids residues), containing catalytic site (the sequences of the enzyme domains according with UniProt Q9NPC4, https://www.uniprot.org/.). The human Gb3/CD77 synthase contains Q at position 211 in contrast to E enzyme with p.Q211E substitution (red dot). Both Gb3/CD77 synthase forms contain two N-glycosylation sites at N_121_ and N_203_ (black dot). Q_full_, fully N-glycosylated of Gb3/CD77 synthase; E_full_, fully N-glycosylated mutein Gb3/CD77 synthase; Q_S123A_, Gb3/CD77 synthase with p.S123A substitution; E_S123A_, mutein Gb3/CD77 synthase with p.S123A substitution; Q_T205A_, Gb3/CD77 synthase with p.T205A substitution; E_T205A_, mutein Gb3/CD77 synthase with p.T205A substitution; Q_S123A_/Q_T205A_, Gb3/CD77 synthase with p.S123A/p.T205A substitutions; E_S123A_/E_T205A_, mutein Gb3/CD77 synthase with p.S123A/p.T205A substitutions [Figure was created with BioRender.com].

In order to check whether the targeted sequons are N-glycosylated, CHO-Lec2 cells expressing the glycovariants were lysed and analyzed by western blotting. The human Gb3/CD77 synthase was detected using the mouse monoclonal anti-A4GALT antibody (clone 5C7). The occupancy of N-glycosylation sites was examined by comparing the electrophoretic mobility of the glycovariants with the fully N-glycosylated enzyme forms (Q_full_ and E_full_). The mobility of single-mutants Q_S123A_, E_S123A_, Q_T205A_ and E_T205A_ was almost the same, and their apparent molecular weight (MW) was lower (about 37 kDa) than their fully N-glycosylated Q_full_ and E_full_ counterparts (39 kDa) (Fig. 2A and Fig. 2B). The double-mutant glycovariants Q_S123A_/Q_T205A_ and E_S123A_/E_T205A_ revealed the lowest apparent MW (32 kDa) of all evaluated proteins. The extra bands recognized by anti-A4GALT antibody (clone 5C7) may have come from proteolytic degradation of the Gb3/CD77 synthase during the cell lysis, whereas the high-molecular weight bands may be enzyme aggregates. In summary, these findings suggest that both N-glycosylation sites of human Gb3/CD77 synthase are occupied (Fig. 2A and Fig. 2B).

**Fig. 2.**
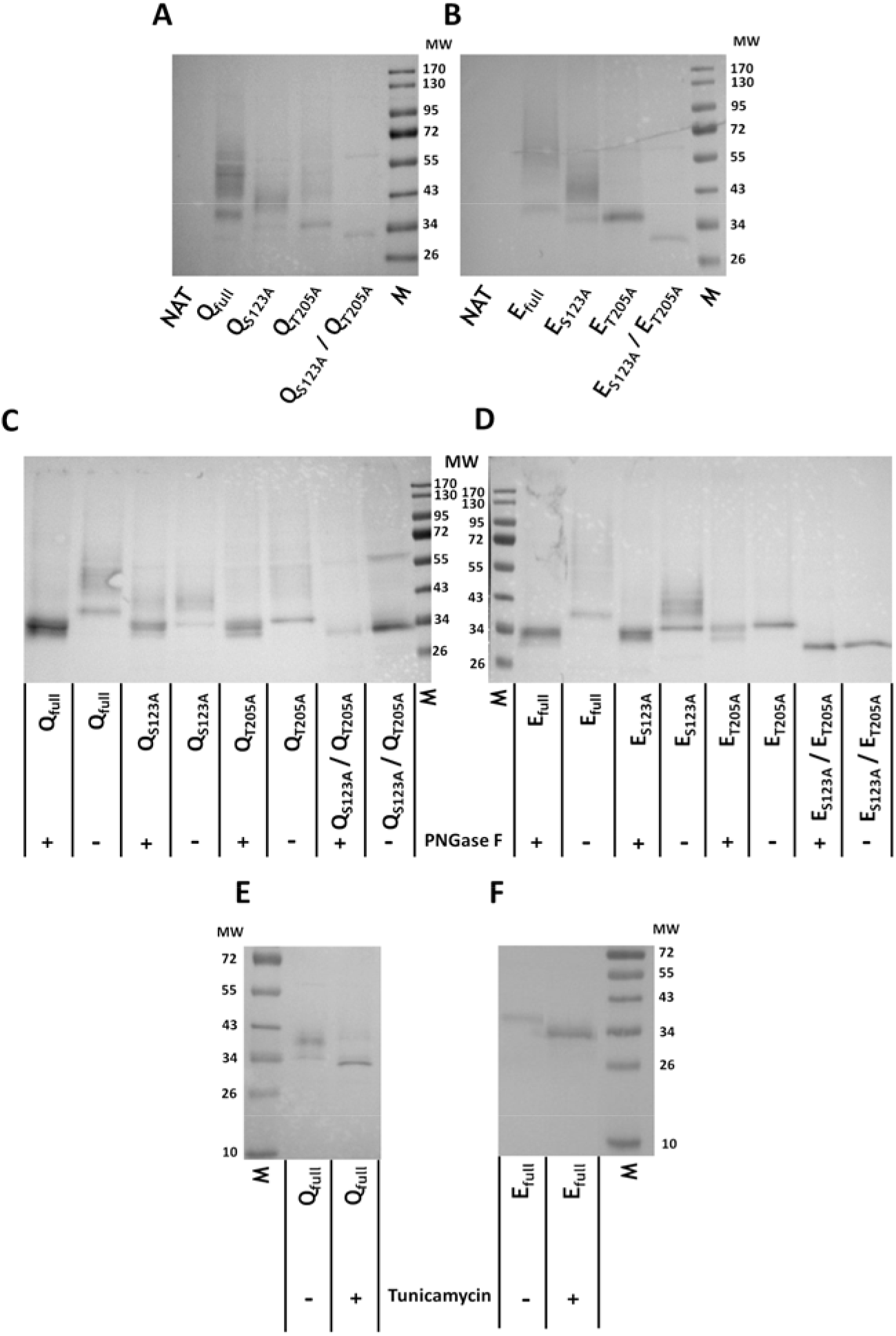
Glycosylation patterns of human Gb3/CD77 synthase and its mutein glycovariants. Western blotting analysis of lysates prepared from non-transfected CHO-Lec2 cells (NAT) and CHO-Lec2 cells expressing fully N-glycosylated enzymes (Q_full_, E_full_) as well as human Gb3/CD77 synthase (Q_S123A_, Q_T205A_, Q_S123A_/Q_T205A_) and its mutein (E_S123A_, E_T205A_, E_S123A_/E_T205A_) glycovariants. Human Gb3/CD77 synthase was stained using anti-A4GALT monoclonal antibody (clone 5C7). Molecular weights of (**A**) human Gb3/CD77 synthase and its (**B**) mutein glycovariants were evaluated. (**C**) Human Gb3/CD77 synthase and its (**D**) mutein glycovariants treated (+) or not (−) with PNGase F. CHO-Lec2 cells transfected with vector encoding (**E**) Q_full_ and (**F**) E_full_ were treated (+) or not (−) with tunicamycin. Molecular weight of the bands are presented as a kDa.

To corroborate these results, CHO-Lec2 cells transfected with vectors encoding the glycovariants and fully N-glycosylated enzymes were lysed and treated with PNGase F, which removes N-glycans from proteins. This generated products whose electrophoretic mobility corresponded with the glycanless glycovariants Q_S123A_/Q_T205A_ and E_S123A_/E_T205A_ (Fig. 2C and Fig. 2D). Additionally, CHO-Lec2 cells expressing the fully N-glycosylated enzymes (Q_full_ and E_full_) were grown in the presence of tunicamycin, which completely inhibits N-glycosylation [Esko J.D., Bertozzi C. et al. 2017]. Both proteins showed similar apparent MW as double-mutants Q_S123A_/Q_T205A_ and E_S123A_/E_T205A_, showing that elimination of N-glycans from these two sites strips the enzyme of all glycans (Fig. 2E and Fig. 2F). In summary, our results indicate that both N-glycosylation sites of human Gb3/CD77 synthase are occupied by N-glycans.

### N_121_ and N_203_ sites have opposing effects on the Gb3/CD77 synthase activity

To examine the activity of human Gb3/CD77 synthase glycovariants in CHO-Lec2 cells, we evaluated the expression of P1PK antigens (Gb3 and P1 for Gb3/CD77 synthase; Gb3, P1 and NOR for its mutein) using flow cytometry (Fig. 3). Two anti-P1 antibodies were used: (1) anti-P1 (clone 650) that reacts with Gb3 and P1 and (2) anti-P1 (clone P3NIL100) that binds only to P1 (Table S1 and Table S2). For detection of the NOR antigen, we used mouse monoclonal nor118 antibody (Table S1 and Table S2) [Duk M., Kusnierz-Alejska G. et al. 2005]. Despite being a quantitative assay, flow cytometry cannot discriminate between GSLs and glycoproteins, nor can it discriminate between Gb3 and P1 when using the anti-P1 antibody (clone 650). CHO-Lec2 cells expressing glycovariants showed different quantities of Gb3, P1 and NOR antigens (determined by estimating antibody binding capacities) (Fig. 4A and Fig. 4B). The highest antibody binding capacities (ABCs) for anti-P1 (650) antibody were found for Q_S123A_ and E_full_ clones, while anti-P1 (P3NIL100) showed highest ABC for Q_full_ and E_full_. The ABC values for anti-P1 (650 and P3NIL100) were the lowest for the double-mutant glycovariants Q_S123A_/Q_T205A_ and E_S123A_/E_T205A_, and partially reduced for E_T205A_ and, to a lesser extent, Q_T205A_ (Fig. 4C and Fig. 4D). These results suggest that the human Gb3/CD77 synthase requires the N_203_-linked glycan for enzyme activity while the N_121_-glycan is dispensable and may even curtail the activity.

**Fig. 3.**
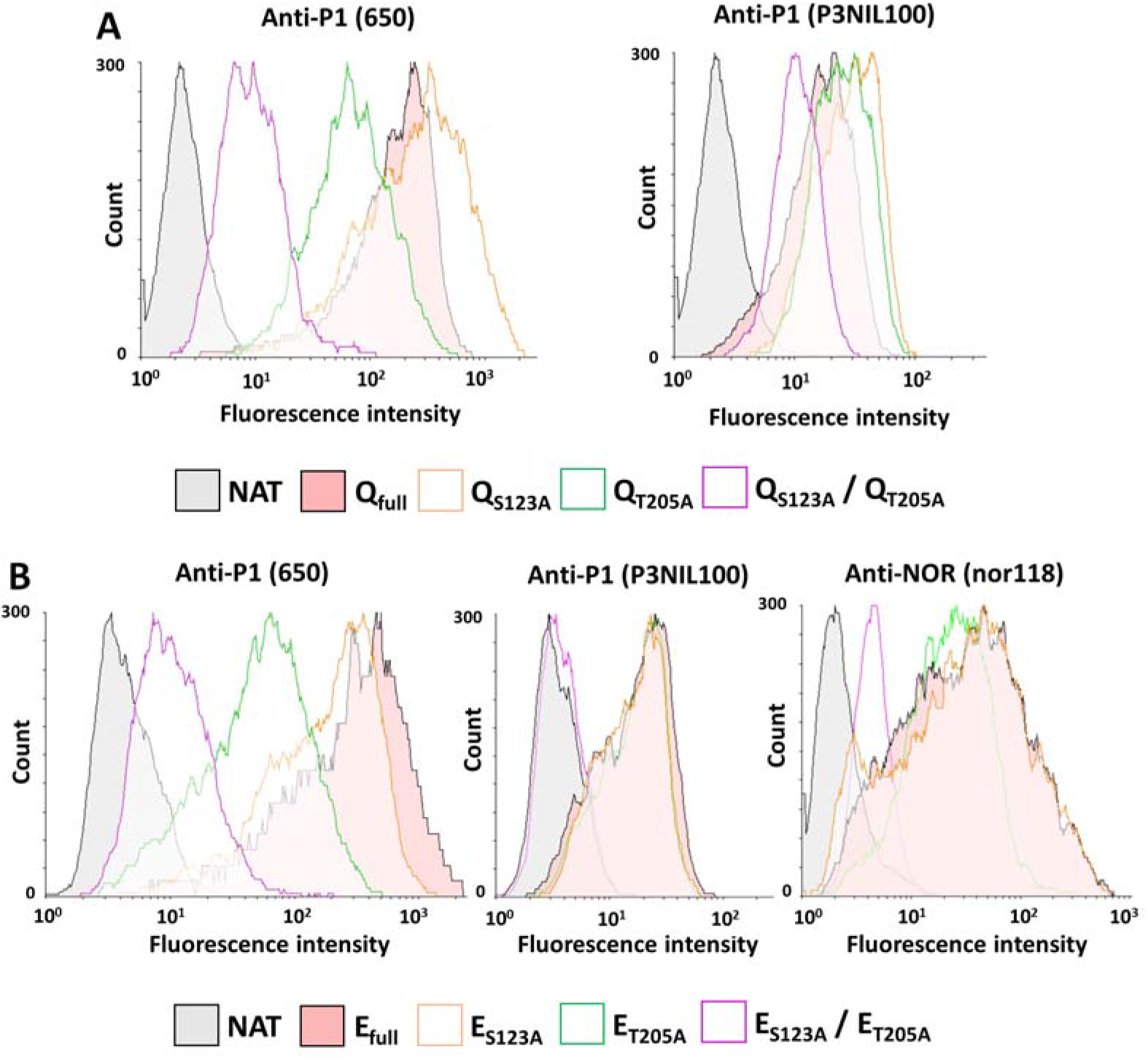
Flow cytometry analysis of CHO-Lec2 cells. The cells were transfected with *A4GALT* genes encoding (**A**) human Gb3/CD77 synthase and its (**B**) mutein glycovariants. The binding of anti-P1 (clones 650 and P3NIL100) and anti-NOR (nor118) antibodies to CHO-Lec2 cells were evaluated. Q_full_, fully N-glycosylated of Gb3/CD77 synthase; E_full_, fully N-glycosylated mutein Gb3/CD77 synthase; Q_S123A_, Gb3/CD77 synthase with p.S123A substitution; E_S123A_, mutein Gb3/CD77 synthase with p.S123A substitution; Q_T205A_, Gb3/CD77 synthase with p.T205A substitution; E_T205A_, mutein Gb3/CD77 synthase with p.T205A substitution; Q_S123A_/Q_T205A_, Gb3/CD77 synthase with p.S123A/p.T205A substitutions; E_S123A_/E_T205A_, mutein Gb3/CD77 synthase with p.S123A/p.T205A substitutions.

**Fig. 4.**
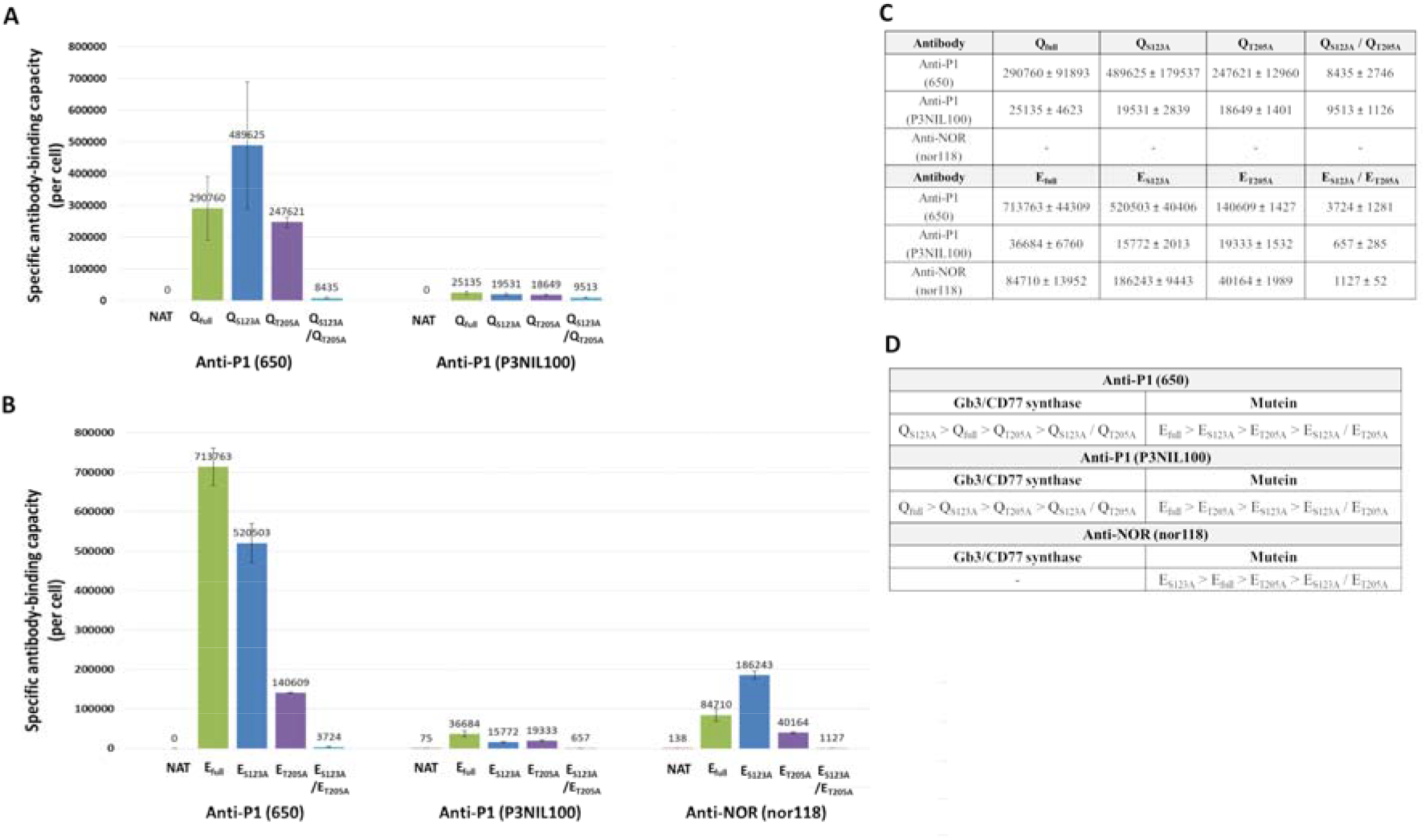
Quantitative flow cytometry of Gb3, P1 and NOR antigens expressing on CHO-Lec2 cells. Antigen binding capacity (ABC) was calculated for anti-P1 (clones 650 and P3NIL100) and anti-NOR (nor118) antibodies per CHO-Lec2 cell. (**A**) Human Gb3/CD77 and its (**B**) mutein glycovariants (median; error bars error bars represent interquartile ranges). (**C**) Antibody binding capacities (ABC) calculated for human Gb3/CD77 and its mutein glycovariants. (**D**) The order of activity level based on the ABC values measured for Gb3/CD77 synthase and its mutein glycovariants. Q_full_, fully N-glycosylated of Gb3/CD77 synthase; E_full_, fully N-glycosylated mutein Gb3/CD77 synthase; Q_S123A_, Gb3/CD77 synthase with p.S123A substitution; E_S123A_, mutein Gb3/CD77 synthase with p.S123A substitution; Q_T205A_, Gb3/CD77 synthase with p.T205A substitution; E_T205A_, mutein Gb3/CD77 synthase with p.T205A substitution; Q_S123A_/Q_T205A_, Gb3/CD77 synthase with p.S123A/p.T205A substitutions; E_S123A_/E_T205A_, mutein Gb3/CD77 synthase with p.S123A/p.T205A substitutions.

To evaluate the activity of human Gb3/CD77 synthase *in vitro*, we used lysates prepared from CHO-Lec2 cells transfected with vectors encoding glycovariants in an enzyme assay using oligosaccharide-polyacrylamide (PAA) conjugates as acceptors. Q_full_ and E_full_ showed the highest activity (Fig. 5A, Fig. 5B and Fig. 5C). Activities of the single-mutant glycovariants were reduced (except for Q_S123A_ with the P1 precursor nLc4-PAA, whose activity was the same as that of Q_full_’s and for E_S123A_ which showed increased activity toward the Gb4-PAA which is a precursor of NOR antigen), while the double-mutant glycovariants Q_S123A_/Q_T205A_ and E_S123A_/E_T205A_ were inactive (Fig. 5A, Fig. 5B and Fig. 5C). Generally, these results suggested that a lack of N-glycan at N_121_ had no significantly effect on the enzyme activity, in contrast to missing the N-glycan at N_203_, when the enzyme shows decreased activity. Finally, the N-glycanless enzymes were completely inactive *in vitro*.

**Fig. 5.**
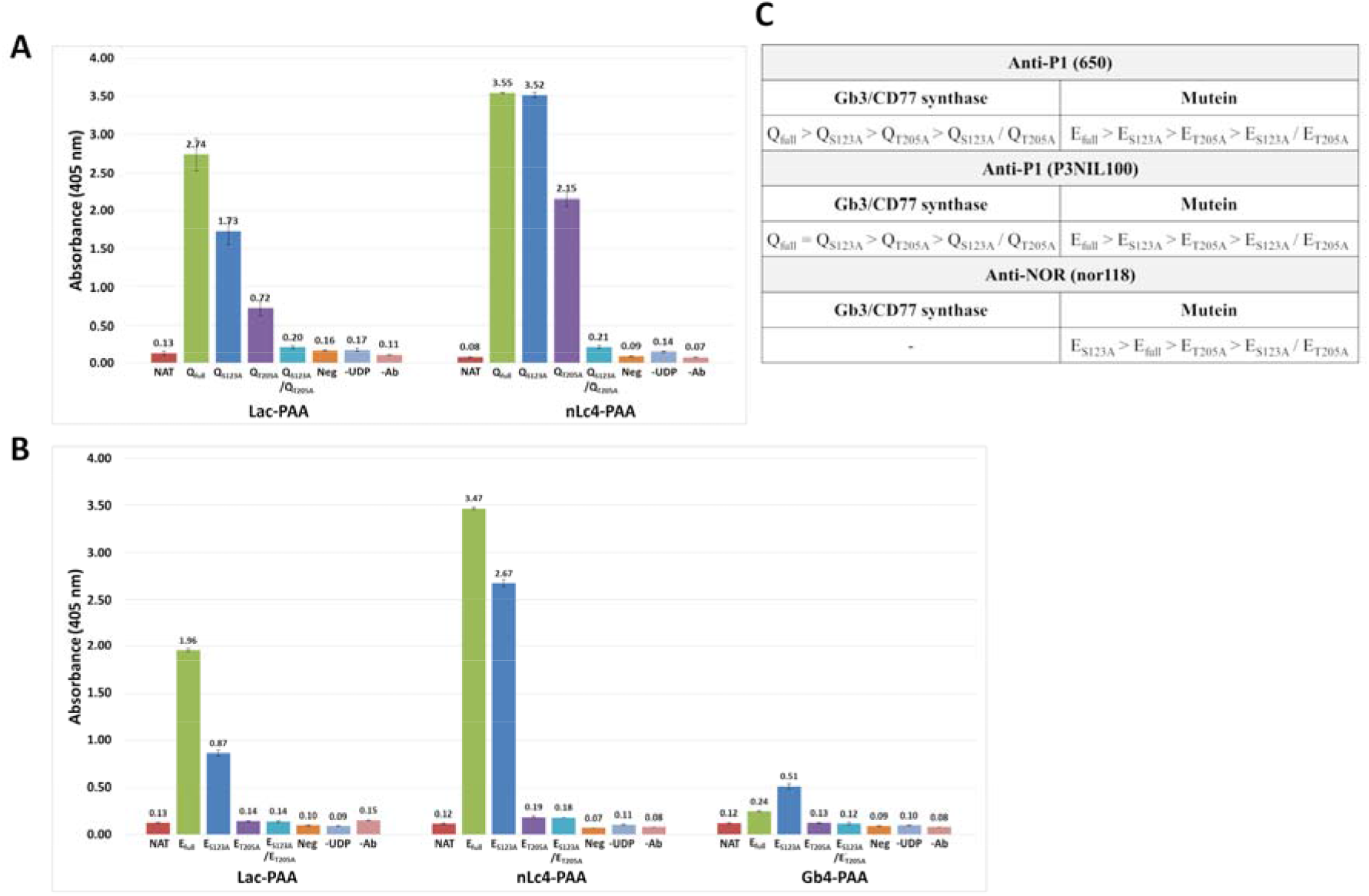
Enzymatic activity of human Gb3/CD77 synthase and its mutein glycovariants in CHO-Lec2 cells lysates. Cell lysates prepared from CHO-Lec2 cells transfected with vectors encoding (**A**) human Gb3/CD77 and its (**B**) mutein glycovariants according to [Cheng C., Guo J. Y. et al. 2016] protocol. (**C**) The order of *in vitro* activity level measured using PAA-conjugates for Gb3/CD77 synthase and its mutein glycovariants. Lac-PAA acceptor is a precursor of Gb3 antigen; nLc4-PAA acceptor is a precursor of P1 antigen; Gb4-PAA acceptor is a precursor of NOR antigen. NAT, non-transfected CHO-Lec2 cells; Q_full_, fully N-glycosylated of Gb3/CD77 synthase; E_full_, fully N-glycosylated mutein Gb3/CD77 synthase; Q_S123A_, Gb3/CD77 synthase with p.S123A substitution; E_S123A_, mutein Gb3/CD77 synthase with p.S123A substitution; Q_T205A_, Gb3/CD77 synthase with p.T205A substitution; E_T205A_, mutein Gb3/CD77 synthase with p.T205A substitution; Q_S123A_/Q_T205A_, Gb3/CD77 synthase with p.S123A/p.T205A substitutions; E_S123A_/E_T205A_, mutein Gb3/CD77 synthase with p.S123A/p.T205A substitutions; Neg, control without lysates containing the enzyme added to reaction; -UDP, control without UDP-Gal donor added to reaction; -Ab, control without added primary antibodies.

Since P1PK GSLs are the main products of human Gb3/CD77 synthase, we analyzed neutral glycosphingolipids isolated from CHO-Lec2 cells transfected with vectors encoding glycovariants with thin-layer chromatography (HPTLC, Fig. 6A and Fig. 6B). In orcinol staining, we found that Gb3Cer and Gb4Cer were the predominant GSLs in all samples. Neither Q- nor E-derived glycovariants produced GSLs detectable with anti-P1 (P3NIL100) antibody (Fig. 6A and Fig. 6B), which suggests that human Gb3/CD77 synthase is unable to synthesize the P1 antigen in CHO-Lec2 cells, most probably due to lack of the acceptor (nLc4).

**Fig. 6.**
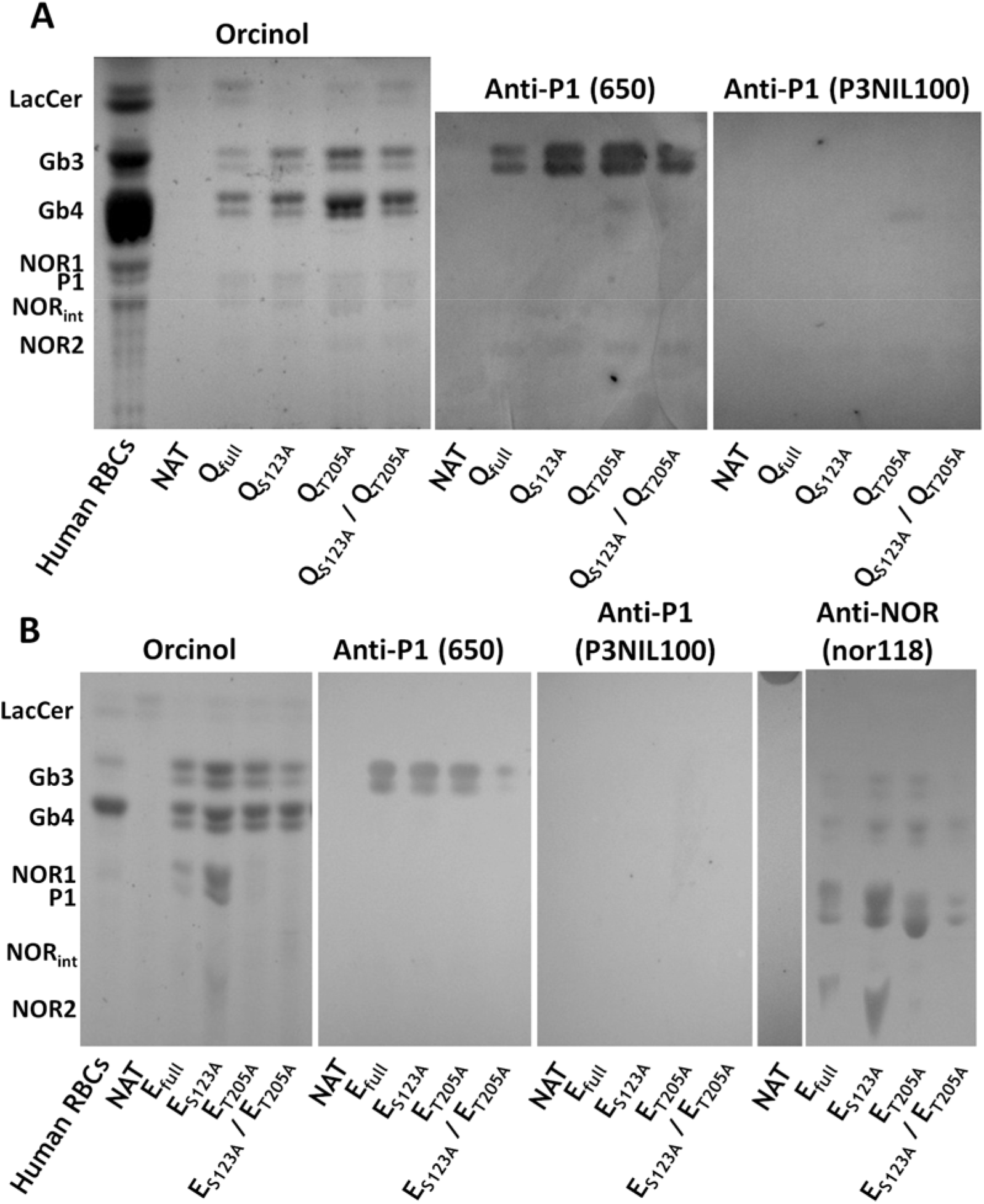
HPTLC analysis of neutral glycosphingolipids extracted from CHO-Lec2 cells. (**A**) Gb3/CD77 synthase and its (**B**) mutein glycovariants analyzed by orcinol staining or by overlaying with anti-P1 (650 and P3NIL100) and anti-NOR (nor118) antibodies. Human RBCs, the GSLs samples of P^1NOR^P^1^ human RBCs; NAT, non-transfected CHO-Lec2 cells; Q_full_, fully N-glycosylated of Gb3/CD77 synthase; E_full_, fully N-glycosylated mutein Gb3/CD77 synthase; Q_S123A_, Gb3/CD77 synthase with p.S123A substitution; E_S123A_, mutein Gb3/CD77 synthase with p.S123A substitution; Q_T205A_, Gb3/CD77 synthase with p.T205A substitution; E_T205A_, mutein Gb3/CD77 synthase with p.T205A substitution; Q_S123A_/Q_T205A_, Gb3/CD77 synthase with p.S123A/p.T205A substitutions; E_S123A_/E_T205A_, mutein Gb3/CD77 synthase with p.S123A/p.T205A substitutions.

The identities of glycosphingolipids derived from CHO-Lec2 transfected cells were confirmed using MALDI-TOF mass spectrometry (Fig. S2 and Fig. S3). The reflectron-positive mode spectrum showed several clusters of ions corresponding to glucosylceramide (GlcCer), lactosylceramide (LacCer) and certain globo-series GSLs (e.g. Gb3, NOR1). The mass differences between the ions (mostly ~ 28 Da, which is the molecular mass of two methylene groups) corresponded to isoforms with ceramides having acyl groups of different lengths (*e.g.* 16:0, 18:0, 20:0, 22:0) but the same long-chain base (d18:1 sphingosine). No P1 antigen-corresponding structures were identified (Fig. S2 and Fig. S3).

Since levels of the P1PK antigens may correlate with the levels of *A4GALT* transcripts in the cells, we examined the expression of genes encoding glycovariants of Q and E enzymes in CHO-Lec2 cells using qPCR. The levels of all transcripts were upregulated in the transfected cells in comparison to the non-transfected CHO-Lec2 (data not shown). The mean threshold cycle (C_T_) revealed no significant differences in transcript levels between cells transfected with different glycovariants (Fig. S4).

There is a general agreement that GSLs are the main acceptors for human Gb3/CD77 synthase, although it was shown recently that the enzyme can also use glycoproteins [Szymczak-Kulus K., Weidler S. et al. 2021; Morimoto K., Suzuki N. et al. 2020; Stenfelt L., Westman J.S. et al 2019]. To find out if the glycovariants differ in acceptor preferences, we analyzed the presence of P1 glycotope on glycoproteins derived from CHO-Lec2 cells transfected with vectors encoding different glycovariants. Using western blotting and anti-P1 antibodies, we found that lysates of cells expressing the Q_full_ and E_S123A_ enzymes revealed the strongest binding (Fig. 7A and Fig. 7B), with the E_S123A_ reaction being markedly stronger than its fully glycosylated counterpart (E_full_). Lysates from the Q_T205A_- and E_T205A_-expressing cells produced weakly recognized glycoproteins, with a slightly stronger reaction in the case of E enzyme (Fig. 7A and Fig. 7B). No binding of anti-P1 (650) antibody was detected in the case of double-mutant glycovariants Q_S123A_/Q_T205A_ and E_S123A_/E_T205A_, although anti-P1 (P3NIL100) weakly recognized a few bands (Fig. 7A and Fig. 7B). These data may suggest that the fully N-glycosylated human Gb3/CD77 synthase and the majority of its glycovariants can efficiently use glycoproteins as acceptors. Moreover, the E_S123A_ enzyme seems to have a higher affinity to glycoprotein acceptors than the other glycovariants.

**Fig. 7.**
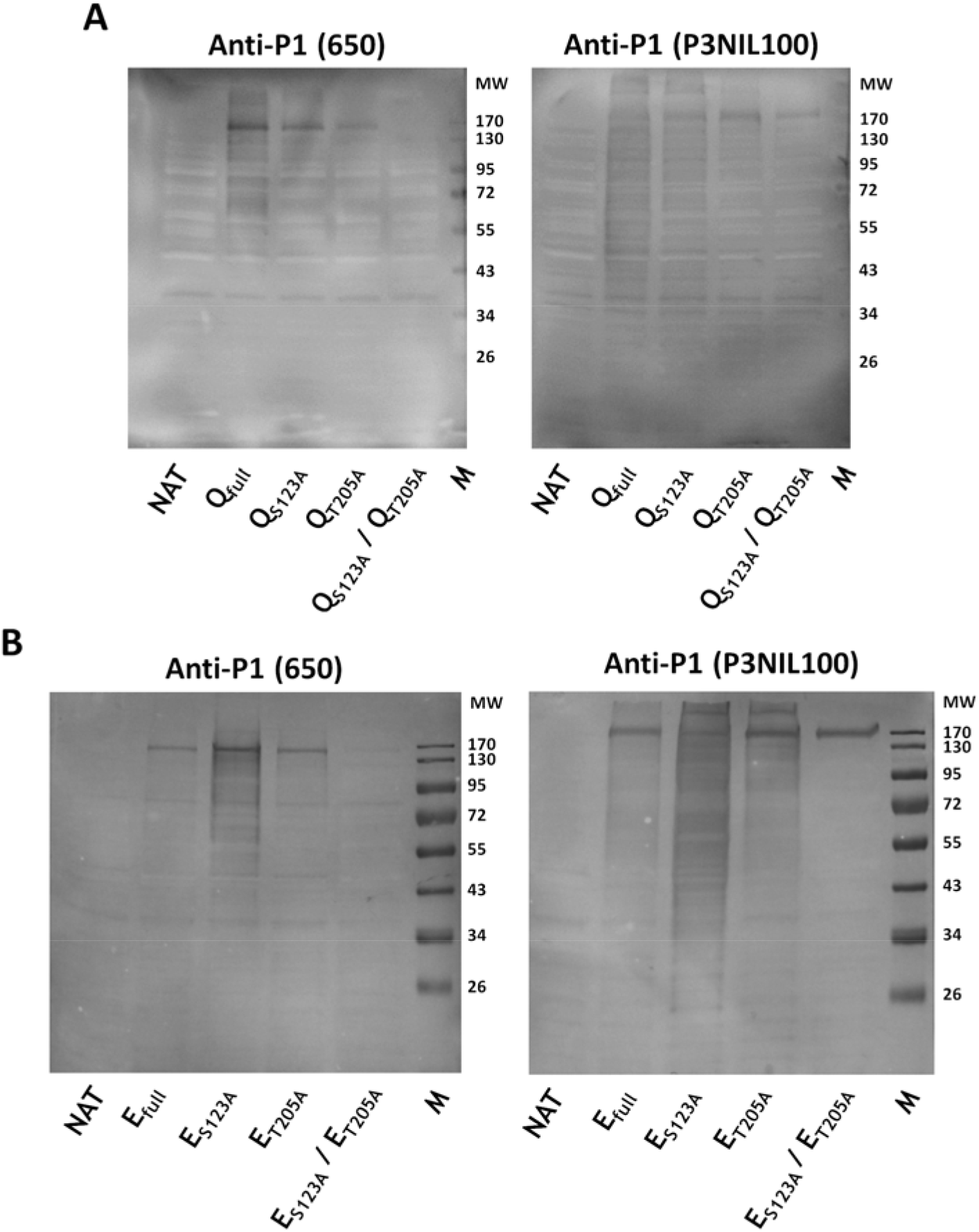
Western blotting analysis of CHO-Lec2 cell lysates stained with anti-P1 (650 and P3NIL100) antibodies. CHO-Lec2 cells transfected with vectors encoding (**A**) Gb3/CD77 synthase and its (**B**) mutein glycovariants. Q_full_, fully N-glycosylated of Gb3/CD77 synthase; E_full_, fully N-glycosylated mutein Gb3/CD77 synthase; Q_S123A_, Gb3/CD77 synthase with p.S123A substitution; E_S123A_, mutein Gb3/CD77 synthase with p.S123A substitution; Q_T205A_, Gb3/CD77 synthase with p.T205A substitution; E_T205A_, mutein Gb3/CD77 synthase with p.T205A substitution; Q_S123A_/Q_T205A_, Gb3/CD77 synthase with p.S123A/p.T205A substitutions; E_S123A_/E_T205A_, mutein Gb3/CD77 synthase with p.S123A/p.T205A substitutions. Molecular weight of the bands are presented as a kDa.

### Cells expressing Gb3/CD77 synthase glycovariants show sensitivity to Shiga toxins

Since the main receptor for Shiga toxins 1 and 2 is Gb3, we evaluated the sensitivity of CHO-Lec2 cells transfected with vectors encoding glycovariants of human Gb3/CD77 synthase and its mutein to these toxins. We found that sensitivity of the cells expressing single-mutant glycovariants of Q and E treated with Stx1 and Stx2 holotoxins was similar or higher than that of the cells expressing fully N-glycosylated enzymes (Fig. 8A and Fig. 8B). The CHO-Lec2 cells expressing double-mutant glycovariants (Q_S123A_/Q_T205A_ and E_S123A_/E_T205A_) were still more sensitive to both Stxs than non-transfected CHO-Lec2 cells, which do not express receptors for Shiga toxins. Viability of Q_S123A_/Q_T205A_ cells was 34% and 35% for Stx1 and Stx2, respectively (Fig. 8A); viability of E_S123A_/E_T205A_ cells was 37% for Stx1 and 39% for Stx2 (Fig. 8B). Cells expressing Q_full_ and single-mutant Q glycovariants were 34-42% and 25-44% viable after treatment with Stx1 and Stx2, respectively; viability of cells expressing E_full_ and the single-mutant E glycovariants was 23-24% and 22-24% upon exposure to Stx1 and Stx2, respectively (Fig. 8A and Fig. 8B). The viability correlated with Gb3 levels on the CHO-Lec2 cells expressing the studied glycovariants. Interestingly, the residual amounts of Gb3 produced by double-mutant glycovariants were sufficient to mediate cytotoxicity.

**Fig. 8.**
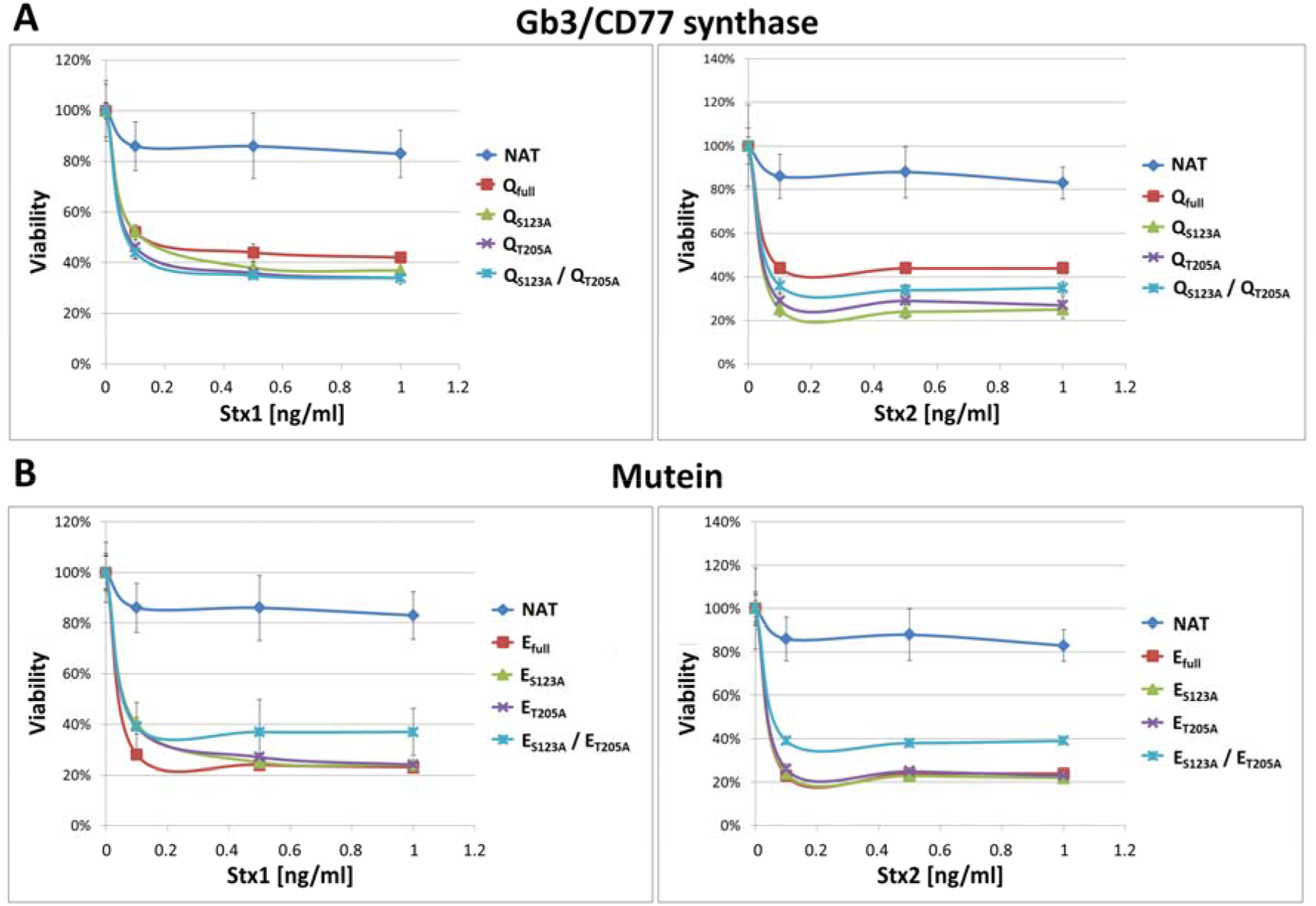
Shiga toxins cytotoxicity analysis. Viability of CHO-Lec2 cells transfected with vectors encoding (**A**) Gb3/CD77 synthase and its (**B**) mutein glycovariants treated with Stx1 and Stx2 holotoxins were evaluated (at least three independent experiments were conducted, each with three technical replicates; error bars are standard deviations; statistical significance when p < 0.05 according to the Kruskal-Wallis ANOVA test). Q_full_, fully N-glycosylated of Gb3/CD77 synthase; E_full_, fully N-glycosylated mutein Gb3/CD77 synthase; Q_S123A_, Gb3/CD77 synthase with p.S123A substitution; E_S123A_, mutein Gb3/CD77 synthase with p.S123A substitution; Q_T205A_, Gb3/CD77 synthase with p.T205A substitution; E_T205A_, mutein Gb3/CD77 synthase with p.T205A substitution; Q_S123A_/Q_T205A_, Gb3/CD77 synthase with p.S123A/p.T205A substitutions; E_S123A_/E_T205A_, mutein Gb3/CD77 synthase with p.S123A/p.T205A substitutions.

### The N_203_ glycan in Gb3/CD77 synthase determines its subcellular localization

To determine whether N-glycosylation affects trafficking and localization of human Gb3/CD77 synthase and its mutein, we evaluated the CHO-Lec2 cells transfected with vectors encoding glycovariants using anti-A4GALT monoclonal antibody (clone 5C7) by immunofluorescence microscopy. It is generally assumed that the human Gb3/CD77 synthase, similarly to other GTs belonging to the glycosyltransferase family 32, is a Golgi-resident enzyme (www.cazy.org); according to Yamaji T. et al. and D’Angelo G. et al., it localizes in *trans*-Golgi network (TGN) [Yamaji T., Sekizuka T. et al. 2019; D’Angelo G., Uemura T. et al. 2013]. Co-immunostaining of Q_full_, E_full_, Q_S123A_, E_S123A_, Q_T205A_ and E_T205A_ glycovariants with anti-A4GALT antibody and anti-syntaxin 16 as a marker of the *trans*-Golgi cisternae revealed that all these enzymes localized in *trans*-Golgi (Fig. 9 and Fig. 10). Q_T205A_, E_T205A_, Q_S123A_/Q_T205A_ and E_S123A_/E_T205A_ glycovariants co-localized with anti-calnexin antibody, which recognizes ER-resident protein (Fig. 9 and Fig. 10). Moreover, the glycovariants Q_T205A_ and E_T205A_ co-localized with both anti-syntaxin 16 and anti-calnexin antibodies, revealing that these enzymes localized both in the *trans*-Golgi and the ER. In contrast, the double-mutant glycovariants Q_S123A_/Q_T205A_ and E_S123A_/E_T205A_ accumulated in the ER only (Fig. 9 and Fig. 10). The residual signals derived from anti-LAMP1 (marker of lysosomes) and anti-A4GALT antibodies were found only in the case of Q_S123A_/Q_T205A_ glycovariant (Fig. 9). Thus, it may be concluded that the fully N-glycosylated enzymes (Q_full_ and E_full_), as well as Q_S123A_ and E_S123A_ glycovariants exit the ER properly and localize in the Golgi. In contrast, the glycovariants with substituted N_203_ site (Q_T205A_ and E_T205A_) partially fail to leave the ER. The glycanless variants (Q_S123A_/Q_T205A_ and E_S123A_/E_T205A_) were found only in the ER. Thus, these findings show that N-glycan at position N_203_ plays a crucial role in trafficking and proper subcellular localization of the human Gb3/CD77 synthase.

**Fig. 9.**
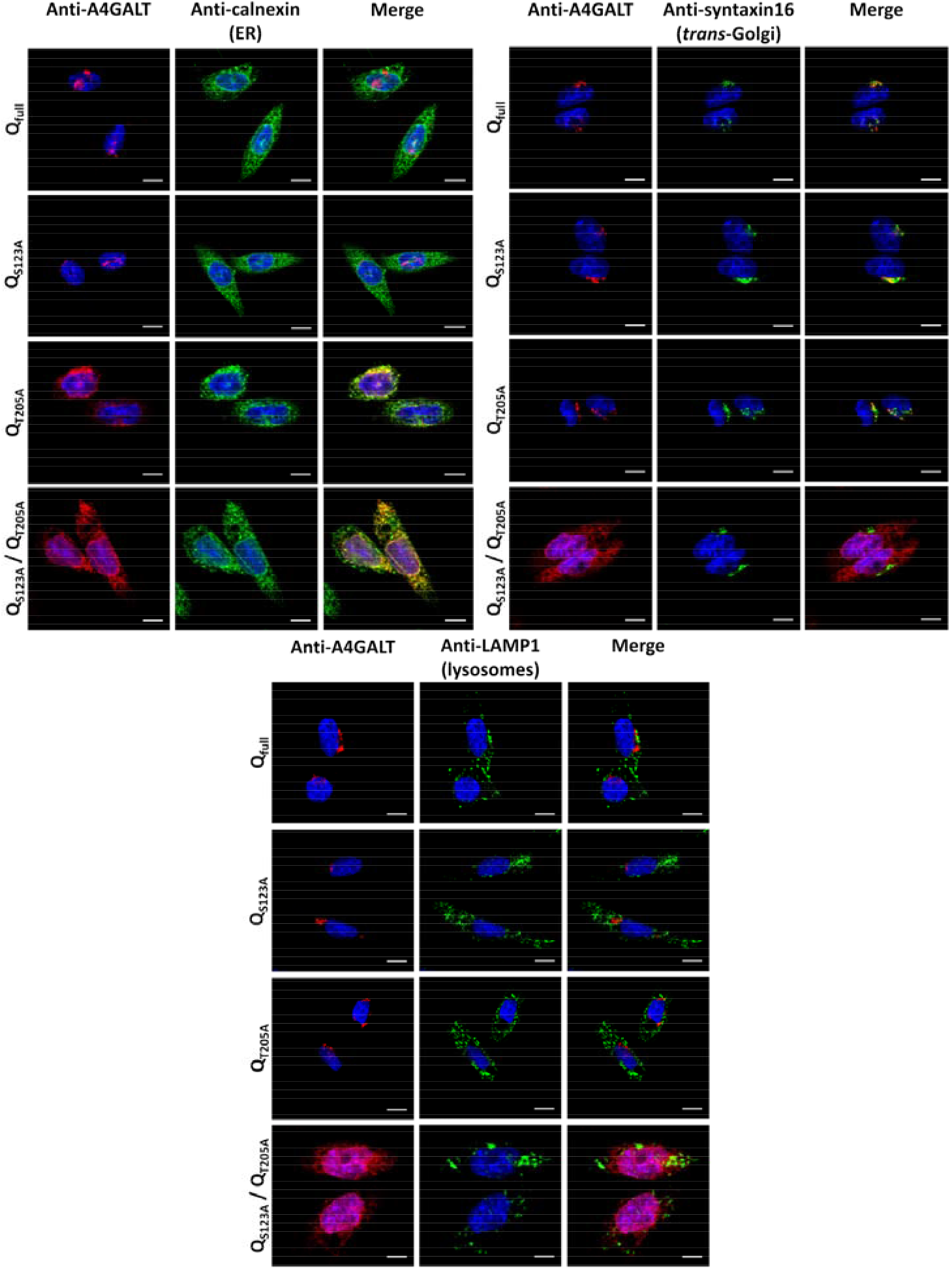
Subcellular localization of human Gb3/CD77 synthase glycovariants in CHO-Lec2 cells using immunofluorescence. The glycovariants were visulized using anti-A4GALT monoclonal antibody (clone 5C7) (red). The cellular organelles, such as Golgi apparatus, endoplasmic reticulum and lysosomes were immunostained by specific antibodies recognizing organellum-specific markers (green). Cell nuclei were counterstained with DAPI (blue). Q_full_, fully N-glycosylated of Gb3/CD77 synthase; Q_S123A_, Gb3/CD77 synthase with p.S123A substitution; Q_T205A_, Gb3/CD77 synthase with p.T205A substitution; Q_S123A_/Q_T205A_, Gb3/CD77 synthase with p.S123A/p.T205A substitutions. Scale bar - 10 μm for fully N-glycosylated and single-mutant glycovariants. Scale bar - 5 μm for double-mutant glycovariants.

**Fig. 10.**
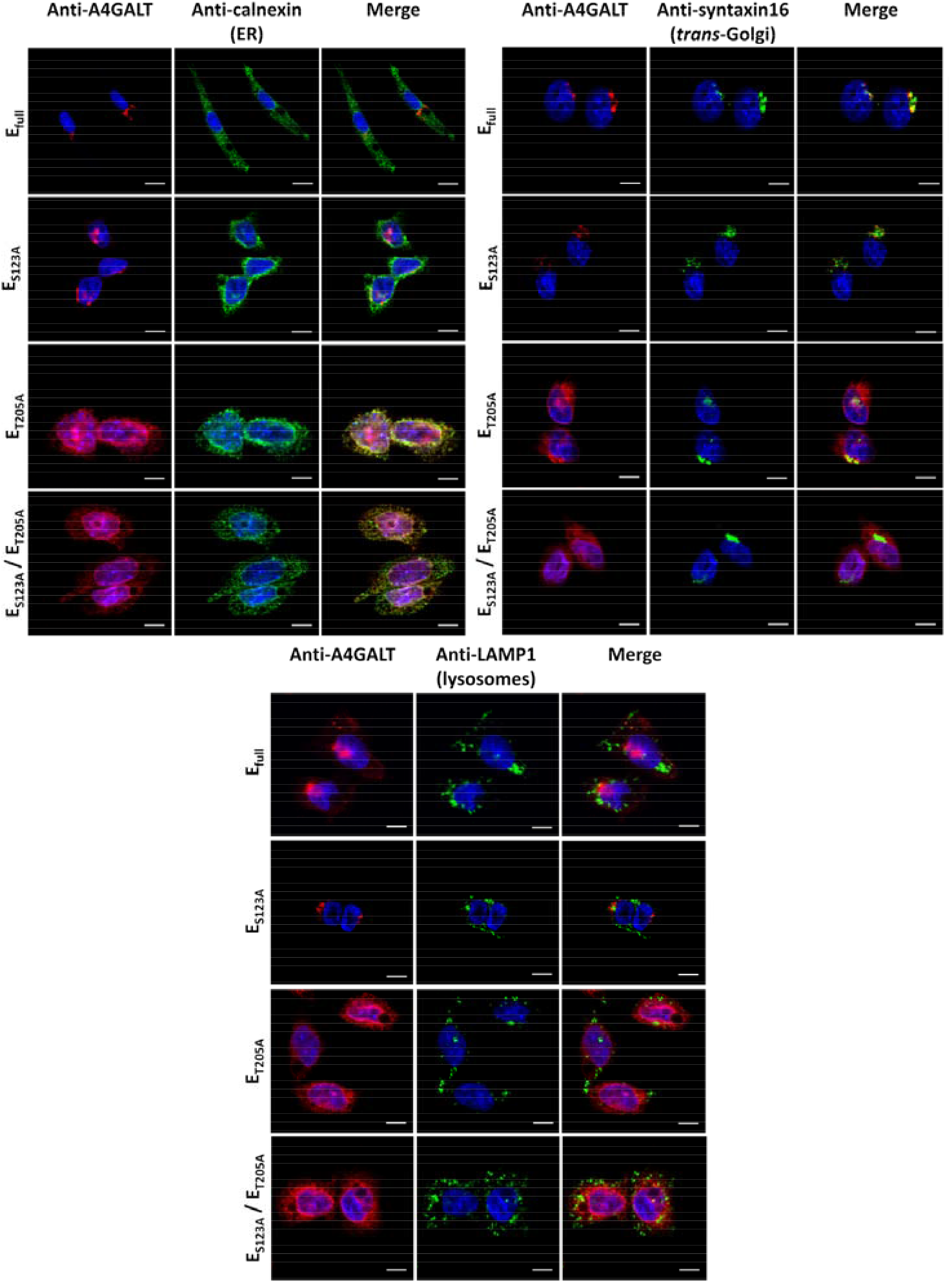
Subcellular localization of mutein glycovariants in CHO-Lec2 cells using immunofluorescence. The glycovariants were visulized using anti-A4GALT monoclonal antibody (clone 5C7) (red). The cellular organelles, such as Golgi apparatus, endoplasmic reticulum and lysosomes were immunostained by specific antibodies recognizing organellum-specific markers (green). Cell nuclei were counterstained with DAPI (blue). E_full_, fully N-glycosylated mutein Gb3/CD77 synthase; E_S123A_, mutein Gb3/CD77 synthase with p.S123A substitution; E_T205A_, mutein Gb3/CD77 synthase with p.T205A substitution; E_S123A_/E_T205A_, mutein Gb3/CD77 synthase with p.S123A/p.T205A substitutions. Scale bar - 10 μm for fully N-glycosylated and single-mutant glycovariants. Scale bar - 5 μm for double-mutant glycovariants.

All glycovariants of human Gb3/CD77 synthase were further evaluated using immunogold reaction with anti-A4GALT monoclonal antibody (clone 5C7). We found that the glycovariants localized in the ER and/or Golgi (Fig. S5 and Fig. S6). In addition, we observed increased numbers of gold nanoparticles corresponding to fully N-glycosylated Q_full_ and E_full_ enzymes as well as Q_S123A_ and E_S123A_ glycovariants, when comparing to glycovariants with p.T205A substitution (Fig. S5 and Fig. S6). In contrast, the double-mutant Q_S123A_/Q_T205A_ and E_S123A_/E_T205A_ glycovariants showed decreased relative numbers of gold nanoparticles in comparison to other glycovariants (Fig. S5 and Fig. S6). These data suggest that glycovariants missing the N_203_-linked glycan were synthesized less efficiently, in contrast to the N_121_ single-mutants, which seemed unaffected by the lack of N-glycan.

## Discussion

N-glycosylation of proteins has important and well-defined functions, but one somewhat overlooked function is regulation of glycosyltransferase activity. The available data on that role were recently reviewed in [Mikolajczyk K., Kaczmarek R. et al. 2020]. Elimination of N-glycans usually does not directly impact catalytic functions, but it may affect the enzyme stability, subcellular localization, secretion and ability to oligomerize, which may influence the enzyme activity. In most cases, altered activity of a non-glycosylated or underglycosylated enzyme is caused by misfolding and accumulation in the ER, which prevents transport to the proper cellular compartment and/or its degradation caused by enhanced aggregation [Mikolajczyk K., Kaczmarek R. et al. 2020; Skropeta D., 2009].

Contrary to the old “one enzyme - one linkage” rule, some GTs are promiscuous [Biswas A., Thattai M., 2020]. One example is human Gb3/CD77 synthase, which can recognize two different disaccharide acceptors, giving rise to Gb3 and the P1 antigen [Kaczmarek R., Duk M. et al. 2016]. A single amino acid substitution p.Q211E, which enables attachment of galactose to another terminal monosaccharide, GalNAc, creating Gal◻1→4GalNAc structure, makes the enzyme even more promiscuous. Extension of acceptor specificity via a single amino acid substitution is a rare phenomenon compared to the more common donor specificity changes. For example, the donor specificity of the ABO transferase depends on two amino acids: the enzyme with c.796C>A (p.L266M) and c.803G>C (p.G268A) substitutions attaches galactose instead of *N*-acetylgalactosamine to the acceptor. Moreover, the cisAB enzyme (with p.L266G substitution) can use either donor substrate, producing both the A and B antigens [Wagner G.K., Pesnot T. et al 2015; Ramakrishnan B., Qasba P.K. et al. 2002].

To date, no studies have fully investigated the influence of N-glycosylation on human Gb3/CD77 synthase activity. Human Gb3/CD77 synthase contains two N-glycosylation sites (N_121_ and N_203_). Previously, we showed that deglycosylated recombinant catalytic domain of Gb3/CD77 synthase expressed in insect cells is inactive (Fig. S1) [Szymczak K., Kaczmarek R. et al. 2016]. That result prompted us to comprehensively evaluate the influence of N-glycosylation on the Gb3/CD77 synthase and its mutein activity using full-length enzyme expressed in CHO-Lec2 cells, which were selected because they do not express an endogenous Gb3/CD77 synthase and are deficient in CMP-sialic acid transporter [Stanley P. 1983; Patnaik S.K., Stanley P. 2006]. Thus, they are incapable of sialylation, leaving a large proportion of acceptors available for α-galactosylation, which otherwise could be consumed by sialyltransferases. In this study, we tried to answer the question whether the two N-glycosylation sites are occupied by N-glycans, and how each of these N-glycans or their lack affects the enzyme activity. To that end, we used site-directed mutagenesis to replace the third amino acid at one or both of the two canonical N-glycosylation sequons with alanine to generate constructs encoding glycovariants [Kaczmarek R., Mikolajewicz K. et al. 2016; Czerwinski M., Kern J. et al. 2007; Leong S.R., Kabakoff R.C. et al. 1994]. We found that elimination of N-glycans from the enzyme affects its enzymatic activity and that disruption of each N-glycosylation site produces different effects. In flow cytometry, the Q_S123A_ glycovariant, which lacks the N-glycan at N_121_, showed similar or increased activity in comparison to the fully N-glycosylated enzyme, but this result was not consistent with *in vitro* activity assays, in which the Q_S123A_ glycovariant revealed decreased activity in comparison with the Q_full_. Moreover, E_S123A_ exhibited reduced activity compared to the E_full_ enzyme in both flow cytometry and *in vitro* assays (although the NOR synthesis was consistently increased). In microscopy, both glycovariants with substituted N_121_ site localized in the Golgi, similarly to the fully N-glycosylated enzymes. Altogether, these findings at the very least show that Gb3/CD77 synthase does not require the N-glycan linked to N_121_ for activity and/or correct subcellular localization, and suggest that it may be more active without it.

In contrast, we found that the N-glycan at position N_203_ in Gb3/CD77 synthase plays an important role in its activity. Flow cytometry and *in vitro* assays revealed reduced quantities of Gb3 and P1 antigen produced by CHO-Lec2 cells expressing the Q_T205A_ and E_T205A_ glycovariants in comparison to the E_full_ enzyme. Immunofluorescence analysis showed that both Q_T205A_ and E_T205A_ localized in two distinct compartments of the cell: in the ER and the *trans*-Golgi. The ER portions of the enzymes were likely inactive. Thus, partial mislocalization in the ER may affect the overall measurable enzyme activity, because only the Golgi-resident enzyme is capable to perform its catalytic functions. Moreover, the decreased enzyme activity of the glycoforms lacking the N_203_ glycan may be associated with reduced enzyme solubility, because the Q_T205A_ and E_T205A_ glycovariants were not detected in culture media (data not shown), suggesting their poor secretion. In addition, such glycovariants did not exhibit any activity in cell lysates, which may suggest that they aggregate. These results suggest that the N-glycan at position N_203_ site plays a crucial role in subcellular localization, solubility and secretion of the enzyme, thus affecting the specific enzyme activity.

The double-mutant glycovariants (Q_S123A_/Q_T205A_ and E_S123A_/E_T205A_) showed only residual activity. The loss of activity was probably caused by altered subcellular localization because both enzymes were found only in the ER. Also, we did not detect any enzymatic activity of these glycovariants in cell lysates, which may have been caused by a decrease in solubility; they were not detected in the culture medium either (data not shown). Overall, these results show that N-glycosylation of human Gb3/CD77 synthase affects its trafficking, solubility and secretion, but the two N-glycans play vastly different roles.

Several studies showed that removal of N-glycans from GTs may cause a decrease in activity (human α1,3/4-fucosyltransferase-III [Christensen L.L., Jensen U.B. et al. 2000]) and/or its complete loss (murine β1,3-galactosyltransferase-IV [Martina J.A., Daniotti J.L. et al. 2000]). The molecular background of activity change may be related to (1) enzyme misfolding, like for human α2,3-sialyltransferase-II [Ruggiero F.M., Vilcaes A.A. et al 2015], (2) altered subcellular localization of the enzyme, *e.g.* human β1,4-galactosyltransferase-IV [Shauchuk A., Szulc B. et al. 2020], (3) changed kinetic parameters, such as rat β1,4-*N*-acetylglucosaminyltransferase-III [Nagai K., Ihara Y. et al 1997], (4) a decrease in the enzyme solubility/secretion, such as human β1,3-*N*-acetylglucosaminyltransferase-II [Kato T., Suzuki M. et al. 2005], (5) enzyme aggregation, such as rat α2,6-sialyltransferase-I [Chen C. and Colley K.J., 2000]. Generally, it may be assumed that enzymes that fail to exit the ER are degraded, so they cannot reveal any activity, as was shown for plant β1,2-xylotransferase from *Arabidopsis thaliana* [Pagny S., Bouissonnie F. et al. 2003]. It should be noted, however, that usually none of these mechanisms contributes to the decrease in enzyme activity independently because many of them intertwine, *e.g.* incorrect localization may be caused by misfolding. In glycoproteins carrying multiple N-glycans, individual chains may vary in importance. In the case of human Gb3/CD77 synthase, its two N-glycans seem to have opposing effects: the N-glycan at position N_121_ seems to be dispensable (with some data, intriguingly, suggesting that its absence may enhance activity and secretion), while the N-glycan at position N_203_ seems to be prerequisite for the enzyme activity, subcellular trafficking and localization. Additionally, N-glycans may impact protein folding and oligomerization, so eliminating N-glycans may change GT activity by altering its ability to oligomerize. Oligomerization may influence subcellular localization in the ER and/or Golgi, and so, indirectly, the activity of a GT [Harrus D., Khoder-Agha F. et al. 2018; Nilsson T., Hoe M.H. et al. 1994]. Previously, we showed that the Q enzyme of human Gb3/CD77 synthase more readily forms homodimers than the E mutein [Kaczmarek R., Suchanowska A. et al. 2013]. It is possible that eliminating N-glycans affects the enzyme’s capability to oligomerize, resulting in changes in activity.

Since Galα1→4Galβ structures on glycosphingolipids and glycoproteins are recognized by Shiga toxins, we examined the presence of glycoprotein products of Gb3/CD77 synthase glycovariants upon their expression in CHO-Lec2 as well as the sensitivity of CHO-Lec2 cells expressing these glycovariants to Shiga toxins. Strikingly, we found that all kinds of the glycovariants were able to synthesize enough receptors to mediate cytotoxic activity of Shiga toxins, even the double-mutant enzymes, which otherwise revealed only residual activity. Notably, HPTLC orcinol staining showed that all CHO-Lec2 clones produced similar amounts of Gb3. This is not entirely unexpected, because HPTLC assays can readily detect and overestimate even small amounts of GSLs. Nevertheless, the amounts of Gb3 produced by double-mutant glycovariants are sufficient to trigger the cytotoxic effects.

In summary, our study shows that human Gb3/CD77 synthase carries two N-glycans, which dramatically differ in importance. The N-glycan at position N_121_ appears to play little to no role in the activity, with some data suggesting that it muzzles the enzyme and limits its secretion. In contrast, the N-glycan at position N_203_ seems to be necessary for activity and correct localization in the Golgi. The glycanless (i.e. double-mutant) variants were stuck in the ER and could not be efficiently secreted.

The dual role of N-glycosylation in the activity of human Gb3/CD77 synthase is intriguing. Attachment of glycans facilitates and is often necessary for the folding of nascent proteins, so a loss of function is an expected corollary of disrupted glycosylation. In contrast, inconsequence or favorability of underglycosylation is counterintuitive. In the case of Gb3/CD77 synthase, the unexpected role of N_121_-linked glycan may have several interesting implications. It may represent a novel regulatory mechanism of preventing potentially detrimental effects of hyperactivity. Hyperactive enzymes may cause serious disorders. One example is the human serine protease coagulation factor IX (FIX), whose hyperfunctional variants are ~5-8-fold more active than the normal FIX and cause hereditary thrombophilias [Simioni P., Tormene D. et al. 2009; Wu W., Xiao L. et al. 2021]. Alternatively, the N_121_-linked glycan may toggle substrate preferences of Gb3/CD77 synthase between glycosphingolipids and glycoproteins. Notably, avian Gb3/CD77 synthases readily use glycoproteins as acceptors, in contrast to the human enzymes, which use mainly glycosphingolipids [Bereznicka A., Modlinska A. et al. 2019]. N-glycosylation of avian Gb3/CD77 synthases has not been studied and perhaps is the missing link in our understanding of this major interspecies difference. In either scenario, mechanistic aspects of the unusual role of N_121_ glycan will require further studies.

## Materials and methods

### Antibodies

Mouse monoclonal anti-A4GALT antibody (clone 5C7) was produced by immunization of C57Bl/6J mice with four subcutaneous injections of the purified human Gb3/CD77 synthase with p.Q211E substitution expressed in insect cells [Kaczmarek R., Duk M. et al. 2016] emulsified in incomplete Freund’s adjuvant (Sigma-Aldrich, St. Louis, MO). Hybridoma secreting antibodies were produced by fusing immune splenocytes with Sp 2.0-Ag14 mouse myeloma using 50% PEG 1500 solution (Sigma-Aldrich, St. Louis, MO) and subcloned by limiting dilution according to standard methodology [Miazek A., Brockhaus M. et al.1997].

### Site-directed mutagenesis

The *A4GALT* gene (NG_007495.2) encoding full-length Gb3/CD77 synthase and its mutein (with p.Q211E substitution) was used for site-directed mutagenesis, as described previously [Kaczmarek R., Mikolajewicz K. et al. 2016]. Briefly, two N-glycosylation sequons N_121_-A-S and N_203_-L-T were disrupted by introducing a codon for alanine in place of serine or threonine, respectively. In the first PCR reaction, two fragments of *A4GALT* gene were created, each containing the overlapping site with an introduced mutation. In the second reaction, the PCR products were duplexed to generate new template DNA. During the overlap extension phase, each fused product was amplified using primers complementary to the pCAG vector (kindly provided by Dr. Peter W. Andrews, University of Sheffield, Sheffield, UK) (PCAG For and PCAG Rev). The resulting full-length gene fragments were digested with XhoI and NotI (Fermentas, Vilnius, Lithuania), cloned into appropriately digested pCAG vector and sequenced (Genomed, Warsaw, Poland) using primers PkSeqFor and PkSeqRev (listed in Table S3). The plasmids were purified using maxi prep kit (Qiagen, Venlo, Netherlands) according to the manufacturer’s instruction. The PCR reaction was performed in a MJ Mini gradient PCR thermal cycler (BioRad, Hercules, CA, USA). 20 μl reaction mixture contained: approximately 200 ng of the template DNA, 0.2 mM forward and reverse primers, 0.2 mM dNTPs, 1.5 mM MgCl_2_, HF polymerase buffer (1:5 dilution), 1 unit Phusion High-Fidelity DNA Polymerase (Fermentas, Vilnius, Lithuania). The DNA fragments were purified with a gel extraction kit (Gel-Out, A&A Biotechnology, Gdynia, Poland). The sequences of primers are shown in Table S3, and the conditions of PCR reactions are shown in Table S4.

### Cell culture and transfection

CHO-Lec2 cells were obtained from the American Type Culture Collection (Rockville, MD, USA). The cells were grown and maintained in a humidified incubator with 5% CO_2_ at 37°C used DMEM/F12 medium (Thermo Fischer Scientific, Inc., Waltham, MA, USA) with 10% fetal bovine serum (Gibco, Inc., Waltham, MA, USA) and Pen-Strep (Gibco, Inc., Waltham, MA, USA). Culture medium was changed every second or third day, and after reaching 85-90% confluence, the cells were subcultured by treatment with trypsin (0.25% trypsin, 137 mM NaCl, 4.3 mM NaHCO_3_, 5.4 mM KCl, 5.6 mM glucose, 0.014 mM phenol red, 0.7 mM EDTA), harvested, centrifuged at 800 x g for 5 min, resuspended in fresh medium and seeded to new tissue culture plates. One day before transfection, cells were seeded at 2 × 10^5^ cells per well in six-well plates, giving at day of transfection about 60% confluence. The medium was replaced with fresh DMEM F12 (without FBS), and after 4 h the cells were transfected using 60 μg polyethyleimine (PEI, Polysciences, Warrington, PA, USA). Plasmid DNA in an amount of 1.5 μg was diluted in buffer containing 0.15 M NaCl and 20 mM HEPES (pH 7.5) and then mixed with PEI. The transfection mixture was incubated for 20 min at room temperature and then added dropwise to each well. After 48 h medium was replaced with fresh DMEM/F12 with 10% FBS and gradient concentration (5, 10, 20 and 50 μg/ml) of puromycin (Sigma-Aldrich, St. Louis, MO). The medium with antibiotic was changed daily for 10 days and then every 2 days. Selection was carried out until the non-transfected control cells were dead, and then cells were sorted using FACS. The cells were harvested using trypsin, washed and suspended in PBS containing 0.5% FBS and 5 mM EDTA at a density of 10^6^ cells/ml. After 1 h incubation at 4°C with anti-P1 (650 and P3NIL100) antibody (1:200 or 1:400, respectively), the cells were washed with PBS containing 0.5% FBS and 1 h incubated with FITC-conjugated goat anti-murine or goat anti-human F(ab’)_2_ antibodies, respectively. Before cell sorting the cells were filtered through a tubes with cell strainer (Falcon® Round-Bottom Tubes with Cell Strainer Cap, Thermo Fisher Scientific, Waltham, MA, USA) to remove cell aggregates. The analysis was carried out using FACS Aria I cell sorter with FACSDiva software (Becton Dickinson).

### Western blotting and lectin blotting

The proteins were separated in the presence of SDS (Sodium Dodecyl Sulfate, Roth, Karlsruhe, Germany) using 10% polyacrylamide gel, according to the Laemmli method [Laemmli U.K., 1970] and visualized with CBB (Coomassie Brilliant Blue R-250, Roth, Karlsruhe, Germany) or transferred to nitrocellulose membrane (Sigma-Aldrich, St. Louis, MO). The PageRuler Prestained Protein Ladder (Thermo Fisher Scientific, Waltham, MA, USA) was used as a protein standard. The proteins fractionated by SDS-PAGE were transferred to the nitrocellulose membrane (Roth) according to the method of Towbin et al. [Towbin H., Staehelin T. et al. 1979] and detected with mouse anti-A4GALT monoclonal antibody (hybridoma supernatant diluted 1:10, clone 5C7) or mouse anti-c-myc (hybridoma supernatant diluted 1:10, clone 9E10) or with biotinylated lectin *Canavalia ensiformis* agglutinin (ConA) (Vector Laboratories, France) in 1 μg/ml in TTBS (0.05% Tween-20/TBS pH 7.5) with 1 mM MgCl_2_, 1 mM MnCl_2_ and 1 mM CaCl_2_. For lectin blotting, the nitrocellulose membrane, before blocking in 5% bovine serum albumin (BSA), was desialylated by treatment with 0.025 M sulfuric acid for 1 h at 80°C.

### PNGase F digestion

Digestion of CHO-Lec2 protein lysates by PNGase F was performed as described [Maszczak-Seneczko D., Olczak T. et al. 2011] in denaturing conditions. Briefly, 1 μl of 10% SDS and 0.7 μl of 1 M DTT was added to 50 μg of transfected or non-transfected CHO-Lec2 cell lysates and the samples were incubated in 95°C for 5 minutes. The deglycosylation reaction was carried out with 500 units of PNGase F for 3 h at 37°C in a final volume of 20 μl. The reaction was stopped with SDS sample buffer and the products were analyzed by immunoblotting. PNGase F digestion in native conditions was carried out without denaturing reagents and skipped incubation at 95°C. Then, the products were used for enzyme activity examination with oligosaccharide-polyacrylamide (PAA) conjugates [Kaczmarek R., Duk M. et al. 2016].

### Enzyme activity evaluation

In order to evaluate enzyme activity in cell lysates, CHO-Lec2 cells transfected with vectors encoding Gb3/CD77 synthase and its mutein with substituted N-glycosylation sites were harvested and lysed according to [Cheng C., Guo J.Y. et al. 2016]. The buffer exchange (from Tris-HCl pH 7.4 to 50 mM sodium cacodylate pH 7.3) was carried out using Amicon Pro 10 kDa cut-off membranes. Activity of de-N-glycosylated recombinant soluble fragment (without transmembrane domain) of human mutein (obtained according to [Kaczmarek R., Duk M. et al. 2016]) was performed under non-denaturing conditions using 500 units of PNGase F for 18 h at 37°C in 500 mM ammonium bicarbonate buffer, pH 7.8. The enzymatic activity was evaluated by ELISA with oligosaccharide-polyacrylamide (PAA) conjugates [Kaczmarek R., Duk M. et al. 2016]. UDP-Gal (Sigma-Aldrich, St. Louis, MO) was used as a donor and three different conjugates were used as acceptors: Galβ1→4Glc-PAA (Lac-PAA, the precursor of Gb3), Galβ1→4GlcNAcβ1→3Galβ1→4Glc-PAA (nLc4-PAA, the precursor of P1), GalNAcβ1→3Galα1→4Galβ1→4Glc-PAA (Gb4-PAA, the precursor of NOR1) [Tuzikov A., Chinarev A. et al. 2021; Bovin N.V., 1998]. ELISA microtiter plates (Nunc, MaxiSorp, Roeskilde, Denmark) were coated overnight in 4°C with conjugates (2 μg/well) in phosphorane buffer (50 mM, pH 7.4). Enzyme samples (100 μg/well) in cacodylate buffer containing Mn^2+^ and the donor substrate (50 mM sodium cacodylate pH 7.4, 14 mM MnCl_2_, 200 μM UDP-Gal, pH 6.3) were loaded in triplicates. The reactions were run overnight (16-18 h) in 37°C. The plates were then washed twice with distilled water and thrice with PBST (137 mM NaCl, 2.7 mM KCl, 10 mM Na_2_HPO_4_, 1.8 mM KH_2_PO_4_, 0.05% Tween 20) and blocked with 5% BSA in PBST. Next, dilutions of antibodies recognizing the reaction products (1:50, 1:100 and 1:100 for anti-P1 P3NIL100, anti-P1 650 and anti-NOR nor118, respectively) were added and incubated for 90 minutes in room temperature, followed by sequential 1-hour incubation with biotinylated anti-mouse or anti-human IgM antibody (each diluted 1:1000) and ExtrAvidin-alkaline phosphatase conjugate (diluted 1:10000). Wash steps were carried out using PBST/1% BSA. Finally, color reactions were developed with *p*-nitrophenyl phosphate (1 mg/ml in Tris-HCl with 1 mM MgCl_2_) (Sigma-Aldrich, St. Louis, MO). Plates were read using 2300 EnSpire Multilabel Reader (PerkinElmer, Waltham, MA) at 405 nm at several time points within 1 hour. Data were analyzed using Microsoft Office Excel (Microsoft Corp, Redmond, WA). Negative controls were set up by adding incomplete reaction mixtures (lacking UDP-Gal or enzyme) to coated wells; by adding complete reaction mixtures to uncoated wells; or by omitting primary or secondary antibodies.

### Flow cytometry

The cells were incubated with 100 μl appropriately diluted primary antibodies (anti-P1 P3NIL100 1:200, anti-P1 650 1:400, anti-NOR nor118 1:20) for 60 min on ice. Then the cells were washed (all washes and dilutions were done with PBS) and incubated with 100 μl (diluted 1:50) FITC-labeled anti-mouse IgM antibody for 40 min on ice in the dark. The cells were washed and approximately 5 × 10^5^ cells were suspended in 750 μl of cold PBS and analyzed by flow cytometry using FACSCalibur (BD Biosciences, Franklin Lakes, NJ, USA). The number of events analyzed was 10000/gated cell population. The analysis of the results was carried out using Flowing software (Perttu Terho, University of Turku, Turku, Finland) [Sahraneshin Samani F., Moore J.K. et al. 2014]. For quantification of cell surface antigens we used Quantum beads (Bio-Rad, Hercules, CA,USA) which carry defined quantities of FITC, and thus allow plotting calibration curves (mean fluorescence intensity versus Molecules of Equivalent Soluble Fluorochrome units). The cells were then analyzed by flow cytometry and the antibody binding capacity (ABC, the number of antibody molecules bound per cell) was calculated by interpolation from the calibration curve as described in the manufacturer’s protocol and based on the fluorophore-to-protein molar ratios of the FITC-antibody conjugates. Negative control results (secondary antibody only) were subtracted from the sample results to obtain specific antibody binding capacities.

### Extraction and purification of glycosphingolipids from CHO-Lec2

The isolation and fractionation of glycosphingolipids and the orcinol staining were performed as described previously [Duk M., Reinhold B.B. et al. 2001]. Cellular lipids were extracted with chloroform/methanol method from 10^7^ transfected or non-transfected CHO-Lec2 cells. The neutral glycosphingolipids were separated from the phospholipids and gangliosides, purified in peracetylated form, then de-O-acetylated, and desalted. Glycosphingolipid samples were solubilized in chloroform/methanol (2:1, v/v), applied to HPTLC plates (Kieselgel 60, Merck, Darmstadt, Germany), and developed with chloroform/methanol/water (55:45:9, v/v/v). The dried plates were immersed in 0.05% polyisobutylmethacrylate (Sigma-Aldrich, St. Louis, MO) in hexane for 1 min, dried, sprayed with TBS (0.05 M Tris buffer, 0.15 M NaCl (pH 7.4), and blocked in 5% HSA. For antibody assays, the plates were successively overlaid with 1) primary antibody diluted in TBS/1% BSA (TBS-BSA) for 1-1.5 h; 2) biotinylated goat anti-mouse Ig antibody (Dako, Glostrup, Denmark), diluted 1:5000 with TBS-BSA; 3) ExtrAvidin-alkaline phosphatase conjugate (Sigma-Aldrich, St. Louis, MO, USA) diluted 1:1000 with TBS/BSA/0.2% Tween 20 for 1 h; and 4) the substrate solution (nitro blue tetrazolium/5-bromo-4-chloro-3-indolyl phosphate, Sigma-Aldrich, St. Louis, MO). Other details were as described previously [Duk M., Reinhold B.B. et al. 2001; Duk M., Singh S. et al. 2007]. Each HPTLC experiment was repeated three times (without significant differences between consecutive repetitions) and GSLs samples were solubilized in the same volumes of chloroform/methanol (2:1, v/v).

### HPTLC orcinol staining

Orcinol staining was performed using standard procedures as previously described [Kuśnierz-Alejska G., Duk M. et al. 1999; Duk M., Reinhold B.B. et al. 2001]. Briefly, dried HPTLC plates were sprayed with a solution of orcinol (0.2% w/v) in 3 M aqueous sulfuric acid and incubated in an oven at 110°C for 10 minutes.

### Quantitative analysis of transcripts level

Total RNA from transfected or non-transfected CHO-Lec2 cells was prepared using Universal RNA Purification Kit (Eurx, Gdansk, Poland) and the complementary DNAs (cDNAs) were synthesized using SuperScript III First-Strand Synthesis kit (Life Technologies, Carlsbad, CA, USA) with oligo(dT) primers. Quantitative polymerase chain reaction (qPCR) was performed on 30 ng of cDNA using the 7500 Fast Real-Time PCR System (Life Technologies, Carlsbad, CA, USA), according to the manufacturer’s instruction. The *A4GALT* transcripts were detected with Custom TaqMan Gene Expression Assay. The ORF sequences were chosen in assay design to enable detection of transcripts originating from plasmids. A predesigned TaqMan assay targeting exon 2–3 boundary (Hs00213726_m1; Life Technologies) was also used to ensure equal amount of the endogenous *A4GALT* transcript in transfected and non-transfected cells. The transcript quantities were normalized to hamster *GAPDH* endogenous control (sequence in Table S5B). All samples were run in triplicates. Data were analyzed using Sequence Detection software Version 1.3.1 (Life Technologies, Carlsbad, CA, USA). Target nucleotide sequences are shown in Table S5, while qPCR conditions are in Table S6.

### MALDI-TOF Mass Spectrometry of CHO-Lec2 GSLs

MALDI-TOF mass spectrometry was carried out on a MALDI TOF/TOF ultrafleXtreme™ instrument (BrukerDaltonics, Bremen, Germany). Samples were dissolved in chloroform/methanol (2:1, v/v). Norharmane (9H-Pyrido[3,4-b]indole, Sigma-Aldrich, St. Louis, MO) was used as a matrix (10 mg/ml, chloroform/methanol, 2:1, v/v). Spectra were scanned in the range of *m/z* 700–1600 in the reflectron-positive mode. External calibration was applied using the Peptide Calibration Standard II (BrukerDaltonics, Bremen, Germany).

### Cytotoxicity assay

2 × 10^4^ CHO-Lec2 cells were seeded in 96-well plates (Wuxi NEST Biotechnology Co., Ltd, China) in complete DMEM/F12. After 24 h the medium was replaced by 100 μl/well of serum-free DMEM/F12 containing 0.1, 0.5 and 1 ng/ml of Stx1 or Stx2 holotoxins (all concentrations were run in triplicates). After 20 h of toxin treatment, 20 μl/well of MTS tetrazolium compound (CellTiter 96® AQueous One Solution Assay, Promega, Madison, WI) was added. Plates were incubated in humidified, 5% CO_2_ atmosphere for 2.5 h, then absorbance at 490 nm was recorded on ELISA plate reader. Background absorbance registered at zero cells/well was subtracted from the data and the absorbance of wells incubated in the medium without Stx was taken as 100% of cell viability. Each experiment was performed at least three times.

### Immunolocalization of Gb3/CD77 synthase

#### Immunogold

For ultrastructural analysis in transmission electron microscopy (TEM) 5 × 10^6^ transfected or non-transfected CHO-Lec2 cells were fixed in cooled 4% formaldehyde solution (FA), diluted in PBS for 20 min at room temperature (RT) (Thermo Fisher Scientific, Waltham, MA, USA). After fixation, the cells were scraped and the cell suspensions were centrifuged 3 times at 2100 x g for 8 minutes followed by rinsing the samples with PBS and distilled water. After adding 1 drop of bovine thrombin (Biomed, Lublin, Poland) to 2 drops of fibrinogen (1mg/ml; Sigma-Aldrich, St. Louis, MO; Merck KGaA) the cells were entrapped within the fibrin clots. Next, the cell clots were post-fixed for 7 min in 0.25% (w/v) osmium tetroxide OsO_4_ diluted in PBS (Serva Electrophoresis, Heidelberg, Germany). Subsequently, the samples were rinsed with PBS 3 times for 5 min.

The cell clots were dehydrated in the increasing concentration of ethanol, EtOH 50%, 70%, 96%, 99.8% (Stanlab, Lublin, Poland) for 10 min at RT. Afterwards, the samples were incubated for 3 h at RT in the mixture of EtOH and LR White resin (Polysciences, Inc., Warrington, PA, USA) in the following proportions: 2:1, 1:1 and 1:2, respectively. Finally, the samples were embedded in pure resin. Polymerization of the resin blocks was carried out at 55°C for 48 h.

LR White blocks were cut into semithin, 600-nm-thick sections with the Histo Diamond Knife (Diatome, Nidau, Switzerland). The semithin sections were stained with a dye solution (Serva, Alchem, Torun, Poland) and closed with a Euparal mounting agent (Carl Roth, Mannheim, Germany), followed by examination on a light microscope in order to remove excessive resin and select a group of no fewer than 30 cells for TEM documentation. Finally, ultrathin, 70-nm-thick sections were cut with the Ultra 45° Diamond Knife (Diatome), arranged with a loop, and collected onto the dull side of nickel grids (200 mesh, Ted Pella, Redding, Ca, USA). Resin-embedded materials were prepared using an ultramicrotome Power Tome XL (RMC, Tucson, USA).

All incubation steps were performed on top of drops of appropriate reagents at RT. First, the ultrathin sections were incubated in 0.02 M glycine (Biotechnology grade, BioShop Canada Inc., Burlington, Canada), dissolved in PBS (1 time for 10 min) to quench free aldehyde groups, followed by gentle rinsing with PBS. Then, the cells were permeabilized 2 times for 5 min with 0.1% Triton X-100 (Reagent grade, BioShop), followed by washing 3 times for 5 min with PBS. In order to block non-specific antigen-binding sites, the grids were transferred for 1 h to 1% bovine serum albumin PBS solution (Albumin fraction V, Carl Roth) and rinsed with PBS for 5 min. For immunogold reaction, the grids were incubated with mouse anti-A4GALT monoclonal antibody (hybridoma supernatant diluted 1:10, clone 5C7) for 1 h, followed by washing the sections in PBS 3 times for 5 min. Subsequently, secondary antibody conjugated with colloidal gold nanoAbcam, Cambridge, UK, preadsorbed) in 1% BSA in PBS (dilution 1:10) was applied for 1 h (dark chamber).

Next, the grids were rinsed with PBS and distilled water, 6 times for 5 min. Additionally, the specimens were post-fixed with 1% glutaraldehyde (Serva Electrophoresis, Heidelberg, Germany) diluted in PBS for 5 min, followed by rinsing with distilled water 3 times for 5 min. To improve contrast, the ultrathin sections were counterstained with uranyl acetate (10 min) and lead citrate trihydrate (5 min) (Serva), and then rinsed 3 times in distilled water. The sections were then examined using TEM JEM-1011 (Jeol, Tokyo, Japan) with the accelerating voltage of 80 kV. Digital micrographs were collected using TEM imaging platform iTEM1233 equipped with a Morada Camera (Olympus, Münster, Germany) at magnifications ranging from 4 to 50K.

#### Immunofluorescence

Millicell EZ slides were used for culturing cells prior to staining. Cells were fixed on the second day of culture with 4% paraformaldehyde (PFA) in PBS added onto slides for 5 minutes. Afterwards, cells were washed 3 times for 5 minutes with PBS. To inhibit non-specific binding sites the slides were incubated 1 h with blocking solution (1% (w/v) BSA and 0.1% (w/v) saponin in PBS). After blocking, cells were incubated with primary antibodies (Table S1) for 3 h at 37°C. Washing (3 × 5 min) was carried out with the blocking solution. Subsequently, slides were incubated with secondary antibodies (Table S1) for 1 h at 37°C. Counterstain with 4◻,6-diamidino-2-phenylindole (DAPI, Sigma-Aldrich, St. Louis, MO) for 10 min at room temperature was performed for nuclei visualization. Finally, slides were mounted onto glass coverslips using fluorescence mounting medium (Dako). Slides were examined with a LEICA SP8 confocal microscope and pictures were analyzed using ImageJ software (NIH).

## Acknowledgements

We thank M.Sc. Michal Kulus (Wroclaw Medical University, Department of Ultrastructural Research, Wroclaw, Poland) for statistical analysis of cytotoxicity assay results, and Dr. Maria Duk (Hirszfeld Institute of Immunology and Experimental Therapy, Wroclaw, Poland) for valuable discussions.

## Abbreviations

GSL: glycosphingolipid

Gb3: globotriaosylceramide

RBC: red blood cell

Stx: Shiga toxin

STEC: Shiga toxin-producing *Escherichia coli*

Stx1B: Shiga toxin 1 B subunit

Stx2B: Shiga toxin 2 B subunit

ConA: *Canavalia ensiformis* agglutinin

ER: endoplasmic reticulum

HUS: hemolytic-uremic syndrome

PNGase F: peptide-N-glycosidase F

ABC: antigen binding capacity

nLc4: paragloboside

Gb4: globoside

HPTLC: high performance thin layer chromatography

qPCR: quantitative polymerase chain reaction

MALDI-TOF: matrix-assisted laser desorption/ionization-time of flight.

## Conflict of interest statement

The authors declare that they have no conflicts of interest with the contents of this article.

## Data availability statement

Raw MS data of GSLs analyzed in the paper are available at the GlycoPOST, Announced ID GPST000166 (https://glycopost.glycosmos.org/).

## Funding and additional information

This work was funded by the Ministry of Science and Higher Education of Poland “Diamentowy Grant” 0097/DIA/2017/46.

## Notes

### Competing Interest Statement

The authors have declared no competing interest.

### Summary of Updates

1. The following figures have been modified: - Fig. S1., - Fig. 3. 2. We added the technical informations to the Manuscript according to Reviewer. 3. According with reviewers suggestions, we revised indicated parts of the manuscript.

## References

Agthe M, Garbers Y, Grötzinger J, Garbers C. Two N-Linked Glycans Differentially Control Maturation, Trafficking and Proteolysis, but not Activity of the IL-11 Receptor. Cell Physiol Biochem. 2018;45:2071–2085.

Akiyama H, Ide M, Yamaji T, Mizutani Y, Niimi Y, Mutoh T, Kamiguchi H, Hirabayashi Y. Galabiosylceramide is present in human cerebrospinal fluid. Biochem Biophys Res Commun. 2021;536:73–79.

Albesa-Jové D, Giganti D, Jackson M, Alzari PM, Guerin ME. Structure-function relationships of membrane-associated GT-B glycosyltransferases. Glycobiology. 2014;24:108–24.

Bereznicka A, Modlinska A, Duk M, Kaczmarek R, Szymczak-Kulus K, Mikolajczyk K, Kapczynska K, Wittek P, Park EY, Piasecki T, Czerwinski M. Avian glycosphingolipid antigens as receptors for Shiga toxin. Conference: Glyco25, XXV International Symposium on Glycoconjugates, 25 - 31.08 At: Milan, Italy.

Biswas A, Thattai M. Promiscuity and specificity of eukaryotic glycosyltransferases. Biochem Soc Trans. 2020;48:891–900.

Bohl T, Bai L, Li H. Recent Progress in Structural Studies on the GT-C Superfamily of Protein Glycosyltransferases. Subcell Biochem. 2021; 96:259–271.

Bovin NV. Polyacrylamide-based glycoconjugates as tools in glycobiology. Glycoconj J. 1998; 15: 431–46.

Breton C, Fournel-Gigleux S, Palcic MM. Recent structures, evolution and mechanisms of glycosyltransferases. Curr Opin Struct Biol. 2012; 5:540–9.

Bruyand M, Mariani-Kurkdjian P, Gouali M, de Valk H, King LA, Le Hello S, Bonacorsi S, Loirat C. Hemolytic uremic syndrome due to Shiga toxin-producing Escherichia coli infection. Med Mal Infect. 2018;48:167–174.

Cavada BS, Pinto-Junior VR, Osterne VJS, Nascimento KS. ConA-Like Lectins: High Similarity Proteins as Models to Study Structure/Biological Activities Relationships. Int J Mol Sci. 2018; 20:30.

Chen C, Colley KJ. Minimal structural and glycosylation requirements for ST6Gal I activity and trafficking. Glycobiology. 2000;10:531–83.

Cheng C, Guo JY, Geng F, Wu X, Cheng X, Li Q, Guo D. Analysis of SCAP N-glycosylation and Trafficking in Human Cells. J. Vis. Exp. 2016;117:e54709.

Christensen LL, Jensen UB, Bross P, Orntoft TF. The C-terminal N-glycosylation sites of the human alpha1,3/4-fucosyltransferase III, -V, and -VI (hFucTIII, -V, adn -VI) are necessary for the expression of full enzyme activity. Glycobiology. 2000;10:931–9.

Cody EM, Dixon BP. Hemolytic Uremic Syndrome. Pediatr Clin North Am. 2019;66:235–246.

Cooling L. Blood Groups in Infection and Host Susceptibility. Clin Microbiol Rev. 2015;28:801–70.

Czerwinski M, Kern J, Grodecka M, Paprocka M, Krop-Watorek A, Wasniowska K. Mutational analysis of the N-glycosylation sites of Duffy antigen/receptor for chemokines. Biochem Biophys Res Commun. 2007;356:816–21.

D’Angelo G, Uemura T, Chuang CC, Polishchuk E, Santoro M, Ohvo-Rekilä H, Sato T, Di Tullio G, Varriale A, D’Auria S, Daniele T, Capuani F, Johannes L, Mattjus P, Monti M, Pucci P, Williams RL, Burke JE, Platt FM, Harada A, De Matteis MA. Vesicular and non-vesicular transport feed distinct glycosylation pathways in the Golgi. Nature. 2013;501:116–20.

Duk M, Kusnierz-Alejska G, Korchagina EY, Bovin NV, Bochenek S, Lisowska E. Anti-alpha-galactosyl antibodies recognizing epitopes terminating with alpha1,4-linked galactose: human natural and mouse monoclonal anti-NOR and anti-P1 antibodies. Glycobiology. 2005;15:109–18.

Duk M, Reinhold BB, Reinhold VN, Kusnierz-Alejska G, Lisowska E. Structure of a neutral glycosphingolipid recognized by human antibodies in polyagglutinable erythrocytes from the rare NOR phenotype. J Biol Chem. 2001;276:40574–82.

Duk M, Singh S, Reinhold VN, Krotkiewski H, Kurowska E, Lisowska E. Structures of unique globoside elongation products present in erythrocytes with a rare NOR phenotype. Glycobiology. 2007;17:304–12.

Esko JD, Bertozzi C, Schnaar. RL. 2017. Chemical Tools for Inhibiting Glycosylation. In Varki A, Cummings RD, Esko JD, Stanley P, Hart GW, Aebi M, Darvill AG, Kinoshita T, Packer NH, Prestegard JH, Schnaar RL, Seeberger PH, editors Essentials of Glycobiology. 3^rd^ edition. Cold Spring Harbor (NY): Cold Spring Harbor Laboratory Press; 2015–2017.

Fiedler K, Simons K. The role of N-glycans in the secretory pathway. Cell. 1995;81:309–12.

Furukawa K, Kondo Y, Furukawa K. 2014. UDP-G al:Lactosylceramide Alpha 1,4-Galactosyltransferase (A4GALT). In: Taniguchi N., Honke K., Fukuda M., Narimatsu H.,Yamaguchi Y, Angata T. Handbook of Glycosyltransferases and Related Genes, pp. 141–147. Springer Japan, Tokyo.

Geyer PE, Maak M, Nitsche U, Perl M, Novotny A, Slotta-Huspenina J, Dransart E, Holtorf A, Johannes L, Janssen KP. Gastric Adenocarcinomas Express the Glycosphingolipid Gb3/CD77: Targeting of Gastric Cancer Cells with Shiga Toxin B-Subunit. Mol Cancer Ther. 2016;15:1008–17.

Goettig P. Effects of Glycosylation on the Enzymatic Activity and Mechanisms of Proteases. Int J Mol Sci. 2016;17:1969.

Harrus D, Khoder-Agha F, Peltoniemi M, Hassinen A, Ruddock L, Kellokumpu S, Glumoff T. The dimeric structure of wild-type human glycosyltransferase B4GalT1. PLoS One. 2018; 13: e0205571.

Jacob F, Anugraham M, Pochechueva T, Tse BW, Alam S, Guertler R, Bovin NV, Fedier A, Hacker NF, Huflejt ME, Packer N, Heinzelmann-Schwarz VA. The glycosphingolipid P◻ is an ovarian cancer-associated carbohydrate antigen involved in migration. Br J Cancer. 2014;111:1634–45.

Jayaprakash NG, Surolia A. Role of glycosylation in nucleating protein folding and stability. Biochem J. 2017;474:2333–2347.

Kaczmarek R, Buczkowska A, Mikołajewicz K, Krotkiewski H, Czerwinski M. P1PK, GLOB, and FORS blood group systems and GLOB collection: biochemical and clinical aspects. Do we understand it all yet? Transfus Med Rev. 2014;28:126–36.

Kaczmarek R, Duk M, Szymczak K, Korchagina E, Tyborowska J, Mikolajczyk K, Bovin N, Szewczyk B, Jaskiewicz E, Czerwinski M. Human Gb3/CD77 synthase reveals specificity toward two or four different acceptors depending on amino acid at position 211, creating P(k), P1 and NOR blood group antigens. Biochem Biophys Res Commun. 2016;470:168–174.

Kaczmarek R, Mikolajewicz K, Szymczak K, Duk M, Majorczyk E, Krop-Watorek A, Buczkowska A, Czerwinski M. Evaluation of an amino acid residue critical for the specificity and activity of human Gb3/CD77 synthase. Glycoconj J. 2016;33:963–973.

Kaczmarek R, Suchanowska A, Lisowska E, Czerwinski M. 2013. Gb3/CD77 synthase (◻1,4-galactosyltransferase) and its variant form, NOR-synthase, exist as dimers. Conference: 38th FEBS Congress, St. Petersburg; 07/2012 At: St. Petersburg Volume: 280S1.

Kaczmarek R, Szymczak-Kulus K, Bereźnicka A, Mikołajczyk K, Duk M, Majorczyk E, Krop-Watorek A, Klausa E, Skowrońska J, Michalewska B, Brojer E, Czerwinski M. Single nucleotide polymorphisms in A4GALT spur extra products of the human Gb3/CD77 synthase and underlie the P1PK blood group system. PLoS One. 2018;13:e0196627.

Kato T, Suzuki M, Murata T, Park EY. The effects of N-glycosylation sites and the N-terminal region on the biological function of beta1,3-N-acetylglucosaminyltransferase 2 and its secretion. Biochem Biophys Res Commun. 2005;329:699–705.

Kattke MD, Gosschalk JE, Martinez OE, Kumar G, Gale RT, Cascio D, Sawaya MR, Philips M, Brown ED, Clubb RT. Structure and mechanism of TagA, a novel membrane-associated glycosyltransferase that produces wall teichoic acids in pathogenic bacteria. PLoS Pathog. 2019;15:e1007723.

Kizuka Y, Kitazume S, Taniguchi N. N-glycan and Alzheimer’s disease. Biochim Biophys Acta Gen Subj. 2017;1861:2447–2454.

Kovbasnjuk O, Mourtazina R, Baibakov B, Wang T, Elowsky C, Choti MA, Kane A, Donowitz M. The glycosphingolipid globotriaosylceramide in the metastatic transformation of colon cancer. Proc Natl Acad Sci USA. 2005;102:19087–92.

Kuśnierz-Alejska G, Duk M, Storry JR, Reid ME, Wiecek B, Seyfried H, Lisowska E. NOR polyagglutination and Sta glycophorin in one family: relation of NOR polyagglutination to terminal alpha-galactose residues and abnormal glycolipids. Transfusion. 1999;39:32–8.

Laemmli UK. Cleavage of structural proteins during the assembly of the head of bacteriophage T4. Nature. 1970;227:680–5.

Lee MS, Tesh V. Roles of Shiga Toxins in Immunopathology. Toxins (Basel). 2019; 11, 212.

Leong SR, Kabakoff RC, Hébert CA. Complete mutagenesis of the extracellular domain of interleukin-8 (IL-8) type A receptor identifies charged residues mediating IL-8 binding and signal transduction. J Biol Chem. 1994;269:19343–8.

Lombard V, Golaconda Ramulu H, Drula E, Coutinho PM, Henrissat B. The carbohydrate-active enzymes database (CAZy) in 2013. Nucleic Acids Res. 2014;42(Database issue):D490–5.

Lowenthal MS, Davis KS, Formolo T, Kilpatrick LE, Phinney KW. Identification of Novel N-Glycosylation Sites at Noncanonical Protein Consensus Motifs. J Proteome Res. 2016;15:2087–101.

Majowicz SE, Scallan E, Jones-Bitton A, Sargeant JM, Stapleton J, Angulo FJ, Yeung DH, Kirk MD. Global incidence of human Shiga toxin-producing Escherichia coli infections and deaths: a systematic review and knowledge synthesis. Foodborne Pathog Dis. 2014;11:447–55.

Martina JA, Daniotti JL, Maccioni HJ. GM1 synthase depends on N-glycosylation for enzyme activity and trafficking to the Golgi complex. Neurochem Res. 2000;25:725–31.

Maszczak-Seneczko D, Olczak T, Jakimowicz P, Olczak M. Overexpression of UDP-GlcNAc transporter partially corrects galactosylation defect caused by UDP-Gal transporter mutation. FEBS Lett. 2011;585, 3090–3094.

Miazek A, Brockhaus M, Langen H, Braun A, Kisielow P. Intrathymic education of alpha beta and gamma delta T cells is accompanied by cell surface expression of RNA/DNA helicase [corrected]. Eur J Immunol. 1997;27:3269–82.

Mikolajczyk K, Kaczmarek R, Czerwinski M. How glycosylation affects glycosylation: the role of N-glycans in glycosyltransferase activity. Glycobiology. 2020;30:941–969.

Miller JJ, Kanack AJ, Dahms NM. Progress in the understanding and treatment of Fabry disease. Biochim Biophys Acta Gen Subj. 2020;1864:129437.

Morimoto K, Suzuki N, Tanida I, Kakuta S, Furuta Y, Uchiyama Y, Hanada K, Suzuki Y, Yamaji T. Blood group P1 antigen-bearing glycoproteins are functional but less efficient receptors of Shiga toxin than conventional glycolipid-based receptors. J Biol Chem. 2020;295:9490–9501.

Nagai K, Ihara Y, Wada Y, Taniguchi N. N-glycosylation is requisite for the enzyme activity and Golgi retention of N-acetylglucosaminyltransferase III. Glycobiology. 1997;7:769–76.

Nilsson T, Hoe MH, Slusarewicz P, Rabouille C, Watson R, Hunte F, Watzele G, Berger EG, Warren G. Kin recognition between medial Golgi enzymes in HeLa cells. EMBO J. 1994; 13: 562–74.

Okuda T, Tokuda N, Numata S, Ito M, Ohta M, Kawamura K, Wiels J, Urano T, Tajima O, Furukawa K, Furukawa K. Targeted disruption of Gb3/CD77 synthase gene resulted in the complete deletion of globo-series glycosphingolipids and loss of sensitivity to verotoxins. J Biol Chem. 2006;281:10230–5.

Pagny S, Bouissonnie F, Sarkar M, Follet-Gueye ML, Driouich A, Schachter H, Faye L, Gomord V. Structural requirements for Arabidopsis beta1,2-xylosyltransferase activity and targeting to the Golgi. Plant J. 2003;33:189–203.

Patnaik SK, Stanley P. Lectin-resistant CHO glycosylation mutants. Methods Enzymol. 2006;416:159–82.

Ramakrishnan B, Qasba PK. Structure-based design of beta 1,4-galactosyltransferase I (beta 4Gal-T1) with equally efficient N-acetylgalactosaminyltransferase activity: point mutation broadens beta 4Gal-T1 donor specificity. J Biol Chem. 2002;277:20833–9.

Rosnoblet C, Peanne R, Legrand D, Foulquier F. Glycosylation disorders of membrane trafficking. Glycoconj J. 2013;30:23–31.

Ruggiero FM, Vilcaes AA, Iglesias-Bartolomé R, Daniotti JL. Critical role of evolutionarily conserved glycosylation at Asn211 in the intracellular trafficking and activity of sialyltransferase ST3Gal-II. Biochem J. 2015;469:83–95.

Ryan SO, Cobb BA. Roles for major histocompatibility complex glycosylation in immune function. Semin Immunopathol. 2012;34:425–41.

Sahraneshin Samani F, Moore JK, Khosravani P, Ebrahimi M. Features of free software packages in flow cytometry: a comparison between four non-commercial software sources. Cytotechnology. 2014;66:555–9.

Shauchuk A, Szulc B, Maszczak-Seneczko D, Wiertelak W, Skurska E, Olczak M. N-glycosylation of the human β1,4-galactosyltransferase 4 is crucial for its activity and Golgi localization. Glycoconj J. 2020;37:577–588.

Simioni P, Tormene D, Tognin G, Gavasso S, Bulato C, Iacobelli NP, Finn JD, Spiezia L, Radu C, Arruda VR. X-linked thrombophilia with a mutant factor IX (factor IX Padua). N Engl J Med. 2009;361:1671–5.

Skropeta D. The effect of individual N-glycans on enzyme activity. Bioorg Med Chem. 2009;17:2645–53.

Stanley P, Taniguchi N, Aebi M. N-Glycans. 2017. In: Varki A, Cummings RD, Esko JD, Stanley P, Hart GW, Aebi M, Darvill AG, Kinoshita T, Packer NH, Prestegard JH, Schnaar RL, Seeberger PH, editors. Essentials of Glycobiology [Internet]. 3rd ed. Cold Spring Harbor (NY): Cold Spring Harbor Laboratory Press; 2015–2017. Chapter 9.

Stanley P. Selection of lectin-resistant mutants of animal cells. Methods Enzymol. 1983;96:157–84.

Stenfelt L, Westman JS, Hellberg Å, Olsson ML. The P1 histo-blood group antigen is present on human red blood cell glycoproteins. Transfusion. 2019;59:1108–1117.

Suchanowska A, Kaczmarek R, Duk M, Lukasiewicz J, Smolarek D, Majorczyk E, Jaskiewicz E, Laskowska A, Wasniowska K, Grodecka M, Lisowska E, Czerwinski M. A single point mutation in the gene encoding Gb3/CD77 synthase causes a rare inherited polyagglutination syndrome. J Biol Chem. 2012;287:38220–30. Epub 2012 Sep 10. Erratum in: J Biol Chem. 2013;288:294.

Szymczak-Kulus K, Weidler S, Bereznicka A, Mikolajczyk K, Kaczmarek R, Bednarz B, Zhang T, Urbaniak A, Olczak M, Park EY, Majorczyk E, Kapczynska K, Lukasiewicz J, Wuhrer M, Unverzagt C, Czerwinski M. Human Gb3/CD77 synthase produces P1 glycotope-capped N-glycans, which mediate Shiga toxin 1 but not Shiga toxin 2 cell entry. J Biol Chem. 2021;296:1–18.

Taujale R, Venkat A, Huang LC, Zhou Z, Yeung W, Rasheed KM, Li S, Edison AS, Moremen KW, Kannan N. Deep evolutionary analysis reveals the design principles of fold A glycosyltransferases. Elife. 2020; 9:e54532.

Towbin H, Staehelin T, Gordon J. Electrophoretic transfer of proteins from polyacrylamide gels to nitrocellulose sheets: procedure and some applications. Proc Natl Acad Sci USA. 1979;76:4350–4.

Tuzikov A, Chinarev A, Shilova N, Gordeeva E, Galanina O, Ovchinnikova T, Schaefer M, Bovin N. 40 years of glyco-polyacrylamide in glycobiology. Glycoconj J. 2021;38:89–100.

Vajaria BN, Patel PS. Glycosylation: a hallmark of cancer? Glycoconj J. 2017;34:147–156.

Varki A. Biological roles of glycans. Glycobiology. 2017;27:3–49.

Wagner GK, Pesnot T, Palcic MM, Jørgensen R. Novel UDP-GalNAc Derivative Structures Provide Insight into the Donor Specificity of Human Blood Group Glycosyltransferase. J Biol Chem. 2015;290:31162–72.

Wu W, Xiao L, Wu X, Xie X, Li P, Chen C, Zheng Z, Ai J, Valencia A, Dong B, Ding Q, Dong B, Wang X. Factor IX alteration p.Arg338Gln (FIX Shanghai) potentiates FIX clotting activity and causes thrombosis. Haematologica. 2021;106:264–268.

Yamaji T, Sekizuka T, Tachida Y, Sakuma C, Morimoto K, Kuroda M, Hanada K. A CRISPR Screen Identifies LAPTM4A and TM9SF Proteins as Glycolipid-Regulating Factors. iScience. 2019;11:409–424.

Zhang H, Zhou M, Yang T, Haslam SM, Dell A, Wu H. New Helical Binding Domain Mediates a Glycosyltransferase Activity of a Bifunctional Protein. J Biol Chem. 2016;291:22106–22117.

