## Supplementary data for "Missing the sweet spot: one of the two N-glycans on human Gb3/CD77 synthase is expendable"

**Contents**

**Table S1**. Antibodies used in the study.

**Table S2.** Specificity of anti-P1 (650 and P3NIL100) and anti-NOR (nor118) antibodies used in western blotting and flow cytometry.

**Table S3.** Nucleotide sequences of primers used in site-directed mutagenesis and sequencing. Changed nucleotides in codons are shown in red.

**Table S4.** PCR conditions used in site-directed mutagenesis.

**Table S5. A)** Target nucleotide sequences within *A4GALT* open reading frame used for design of Custom TaqMan Gene Expression Assay. **B)** Target nucleotide sequences within hamster *GAPDH* open reading frame (NM_001244854.2) and TaqMan probe nucleotide sequence used for design of Custom TaqMan Gene Expression Assay (endogenous control).

**Table S6**. qPCR conditions used for quantitative analysis of *A4GALT* transcripts.

**Figure S1.** Activity of human recombinant mutein obtained in insect cells after de-N-glycosylation.

**Figure S2.** Reflectron-positive mode MALDI-TOF mass spectra of glycosphingolipids isolated from CHO-Lec2 cells expressing the Gb3/CD77 synthase glycovariants.

**Figure S3.** Reflectron-positive mode MALDI-TOF mass spectra of glycosphingolipids isolated from CHO-Lec2 cells expressing glycovariants of mutein.

**Figure S4.** Comparisons of mean threshold cycle (C_T_) values between Gb3/CD77 synthase and its mutein glycovariants.

**Figure S5.** Subcellular localization of human Gb3/CD77 synthase glycovariants in CHO-Lec2 cells using immunogold reaction.

**Figure S6.** Subcellular localization of mutein glycovariants in CHO-Lec2 cells using immunogold reaction.

**Table S1**. Antibodies used in the study.

| **Antibody** | **Clonality** | **Dilution IF / WB / HPTLC / FACS** | **Host** | **Manufacturer** |
| --- | --- | --- | --- | --- |
| Anti-A4GALT | Monoclonal,  clone 5C7 | 1:10  (WB) | Mouse | Hybridoma supernatant |
| Anti-c-myc | Monoclonal,  clone 9E10 | 1:10  (WB) | Mouse | Hybridoma (purchased from ATCC) supernatant |
| Anti-6x-His | Monoclonal,  clone HIS.H8 | 1:1000  (WB, HPTLC) | Mouse | Thermo Fischer Scientific |
| Anti-P1 | Monoclonal,  clone 650 | 1:100  (WB, HPTLC, FACS) | Mouse | Ce-Immundiagnostika |
| Anti-P1 | Monoclonal,  clone P3NIL100 | 1:100  (WB, HPTLC, FACS) | Human | Immucor Inc. |
| Anti-NOR | Monoclonal,  clone nor118 | 1:20, 1:100  (WB, HPTLC, FACS) | Mouse | Hybridoma supernatant |
| Biotynylated anti- IgG/A/M (H/L) | Polyclonal | 1:1000  (WB) | Goat | Bio-Rad Laboratories |
| Anti-mouse IgM-FITC | Polyclonal | 1:100  (FACS) | Goat | Thermo Fisher Scientific |
| Anti-human IgM-FITC | Polyclonal | 1:100  (FACS) | Goat | Thermo Fisher Scientific |
| Anti-mouse IgG-FITC | Monoclonal | 1:100  (FACS) | Goat | Santa Cruz Biotechnology |
| Anti-calnexin | Polyclonal | 1:100  (IF) | Rabbit | Abcam |
| Anti-syntaxin16 | Monoclonal | 1:100  (IF) | Rabbit | Abcam |
| Anti-LAMP1 | Polyclonal | 1:100  (IF) | Rabbit | Abcam |
| Anti-rabbit Alexa Fluor 568 (secondary) | Polyclonal | 1:1000 (IF) | Goat | Thermo Fisher Scientific |
| Anti-mouse Alexa Fluor 488 (secondary) | Polyclonal | 1:1000 (IF) | Goat | Thermo Fisher Scientific |

**Table S2.** Specificity of anti-P1 (650 and P3NIL100) and anti-NOR (nor118) antibodies used in western blotting and flow cytometry. Cer, ceramide; R, core N-glycan structure; GSL, glycosphingolipid; GP, glycoprotein.

| **Glycan structure** | **Anti-P1** | | **Anti-NOR**  **(nor118)** |
| --- | --- | --- | --- |
|  | **650** | **P3NIL100** |  |
| **Gb3**  (Galα1→4Galβ1→4Glc-Cer) | **+** | **-** | **-** |
| **P1 on GSL**  (Galα1→4Galβ1→4GlcNAcβ1→3Galβ1→4Glc-Cer) | **+** | **+** | **-** |
| **P1 on GP**  (Galα1→4Galβ1→4GlcNAcβ1→R) | **+** | **+** | **-** |
| **NOR1**  (Galα1→4GalNAcβ1→3Galα1→4Galβ1→4Glc-Cer)  **NOR2** (Galα1→4GalNAcβ1→3Galα1→4GalNAcβ1→3Galα1→4Galβ1→4Glc-Cer) | **-** | **-** | **+** |

**Table S3.** Nucleotide sequences of primers used in site-directed mutagenesis and sequencing. Changed nucleotides in codons are shown in red.

| **Primers** | **Sequence [5' → 3']** |
| --- | --- |
| PkSeqFor | TCGCACTCATGTGGAAG |
| PkSeqRev | AGTACATTTTCATGGCCT |
| Pkcon_S123A_sens | GC AAC GCC GCA CTG CCC CGG CAC |
| pCAGanty | ACA AAC GCA CAC CGG CCT TAT TCC |
| Pkcon_S123A_anty | GTG CCG GGG CAG TGC GGC GTT GC |
| pCAGsens | CGT GCT GGT TGT TGT GCT GTC TCA |
| Pkcon_T205A_sens | CTG CGG AAC CTG GCA AAC GTG CTG G |
| PCAGanty | ACA AAC GCA CAC CGG CCT TAT TCC |
| Pkcon_T205A_anty | C CAG CAC GTT TGC CAG GTT CCG CAG |
| pCAGsens | CGT GCT GGT TGT TGT GCT GTC TCA |

**Table S4.** PCR conditions used in site-directed mutagenesis.

| **Mutagenesis – first step** | | | | **Mutagenesis – second step** | | |
| --- | --- | --- | --- | --- | --- | --- |
|  | **Temp. [°C]** | **Time [s]** | **Cycle** | **Temp. [°C]** | **Time [s]** | **Cycle** |
| Initial denaturation | **94** | **180** | **1** | **94** | **180** | **1** |
| Denaturation | **94** | **15** | **30** | **94** | **15** | **30** |
| Annealing | **62-72** | **20** | **30** | **70** | **20** | **30** |
| Extension | **72** | **10** | **30** | **72** | **20** | **30** |
| Final extension | **72** | **300** | **1** | **72** | **300** | **1** |

**Table S5. A)** Target nucleotide sequences within *A4GALT* open reading frame used for design of Custom TaqMan Gene Expression Assay. **B)** Target nucleotide sequences within hamster *GAPDH* open reading frame (NM_001244854.2) and TaqMan probe nucleotide sequence used for design of Custom TaqMan Gene Expression Assay (endogenous control).

**A)**

| **Name of target sequence** | **Sequence [5' → 3']** |
| --- | --- |
| A4gtf | CTGCACCCT |
| A4gtr | TTCTCAAGAAC |

B)

| **Name of target sequence** | **Sequence [5' → 3']** |
| --- | --- |
| Gapdhf | TGGAAAGCTTGTCATCAAC |
| Gapdhr | GAAGACGCCAGTAGATTCC |

| **TaqMan probe** | **Sequence [5' → 3']** |
| --- | --- |
| GAPDH | AGGCCATCACCATCTTCCAG |

**Table S6**. qPCR conditions used for quantitative analysis of *A4GALT* transcripts.

| qPCR system | Reaction format | Reaction volume | Thermal cycling conditions | | | |
| --- | --- | --- | --- | --- | --- | --- |
|  |  | | Parameter | Initial denaturations | PCR (40 cycles) | |
|  |  |  |  |  | Denaturation | Annealing/Extension |
|  |  |  | Temperature [°C] | 95 °C | 95 °C | 60 °C |
| 7500 Fast | 96-well plate | 20 µl | Time (mm:ss) | 10:00 | 0:15 | 1:00 |

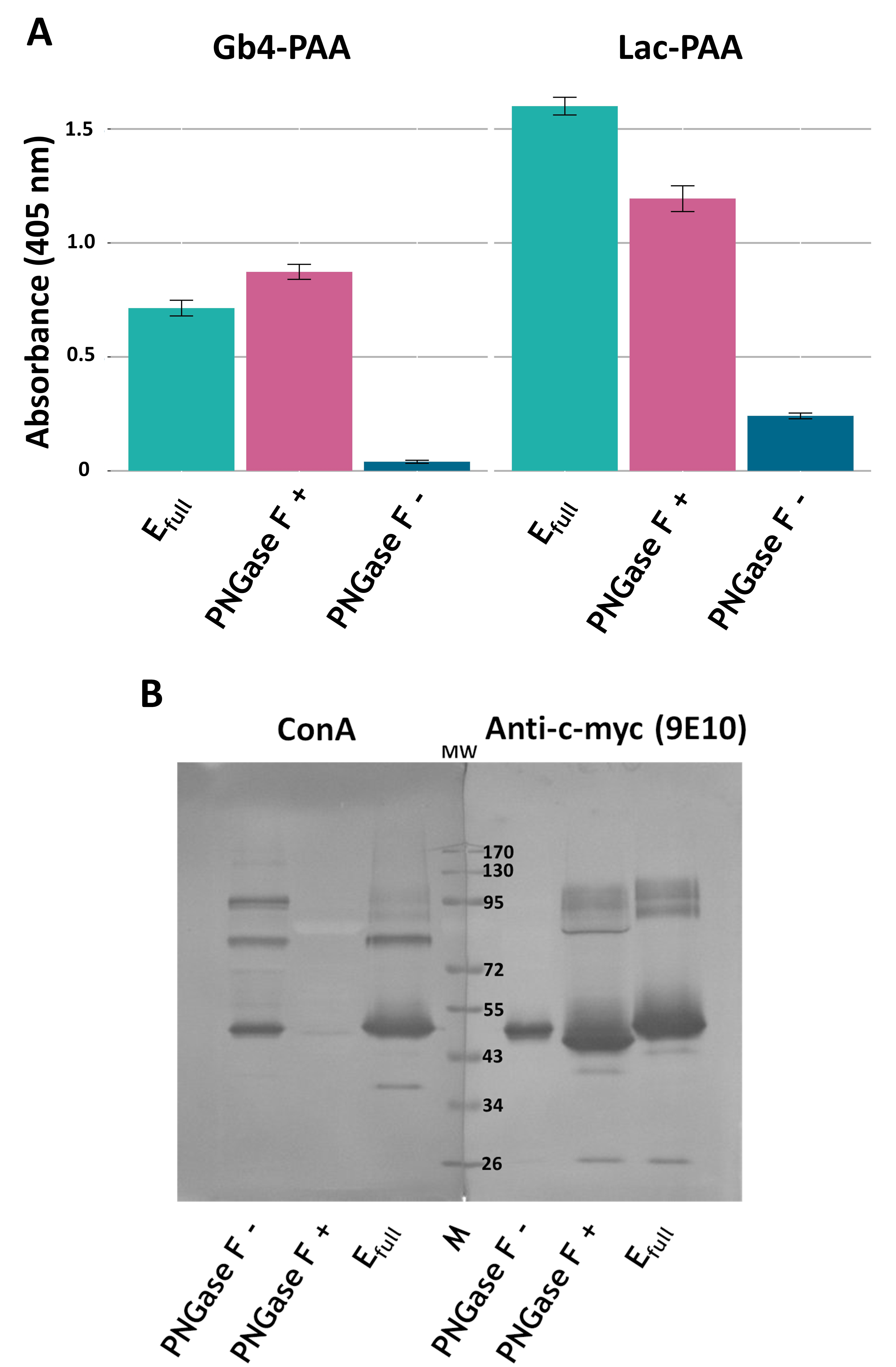

Fig. S1. **Activity of human recombinant mutein obtained in insect cells after de-N-glycosylation.** (**A**) *In vitro* activity of untreated human mutein (E_full_), enzyme treated with PNGase F (PNGase F +) or incubated in deglycosylation buffer (PNGase F -). Enzymatic activity was evaluated using PAA-conjugates, serving as precursors of Gb3 antigen (Lac-PAA acceptor) and NOR (Gb4-PAA acceptor) [Szymczak K., Kaczmarek R. et al. 2016]. (**B**) Lectin blotting and western blotting of untreated human mutein (E_full_), enzyme treated with PNGase F (PNGase F +) and treated only with deglycosylation buffer (PNGase F -). The blots were overlaided with ConA lectin, which is specific for core oligosaccharide of N-glycans with α-linked mannose [Cavada B.S., Pinto-Junior V.R. et al. 2018] and with anti-c-myc antibody (clone 9E10), which recognizes c-myc tag at C-terminus of soluble human Gb3/CD77 synthase [Kaczmarek R., Duk M. et al. 2016].

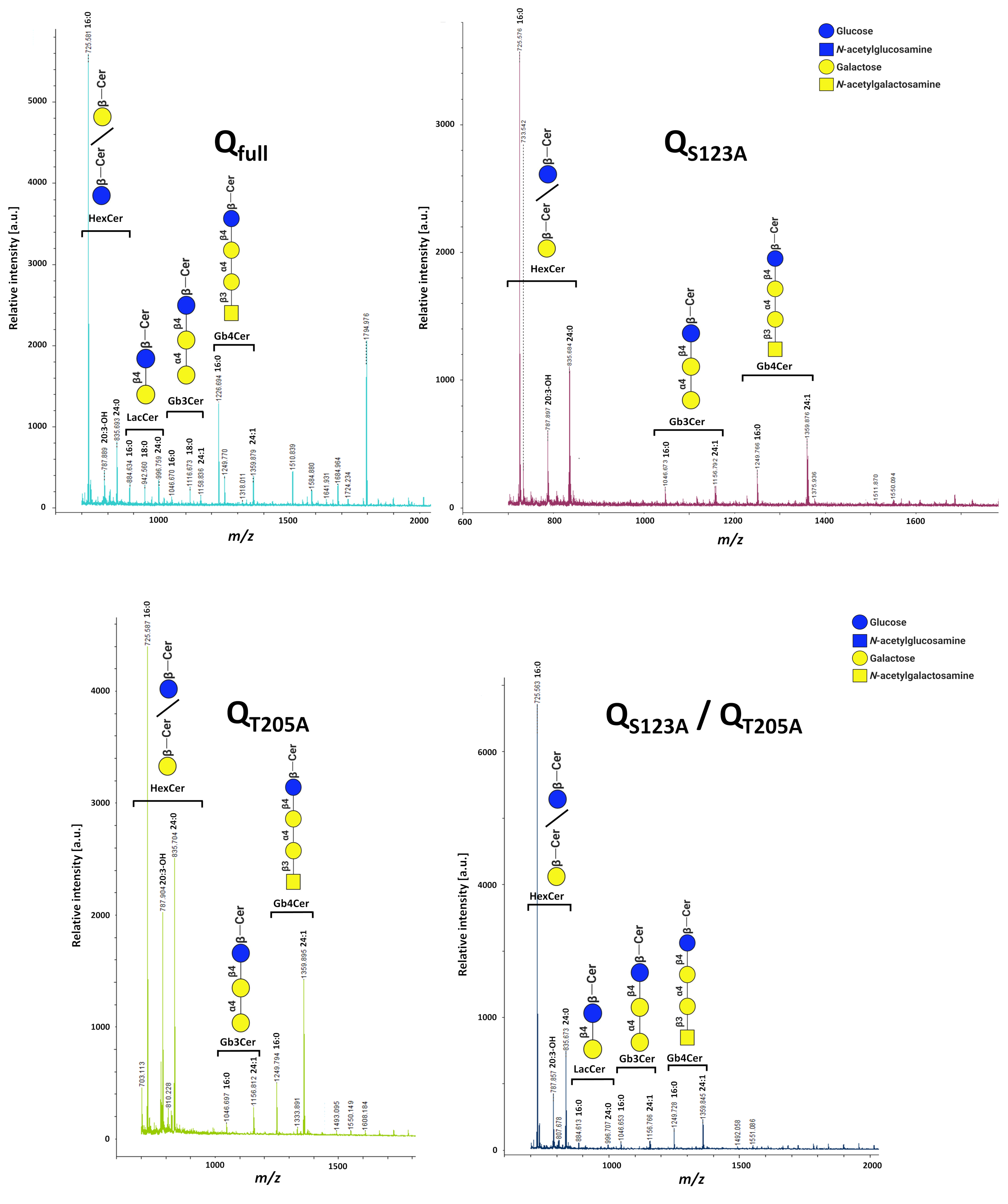

Fig. S2. **Reflectron-positive mode MALDI-TOF mass spectra of glycosphingolipids isolated from CHO-Lec2 cells expressing the Gb3/CD77 synthase glycovariants.** The GSLs samples from CHO-Lec2 cells transfected with vectors encoding fully N-glycosylated Q_full_ enzyme as well as Q_S123A_, Q_T205A_ and Q_S123A_/Q_T205A_ glycovariants.

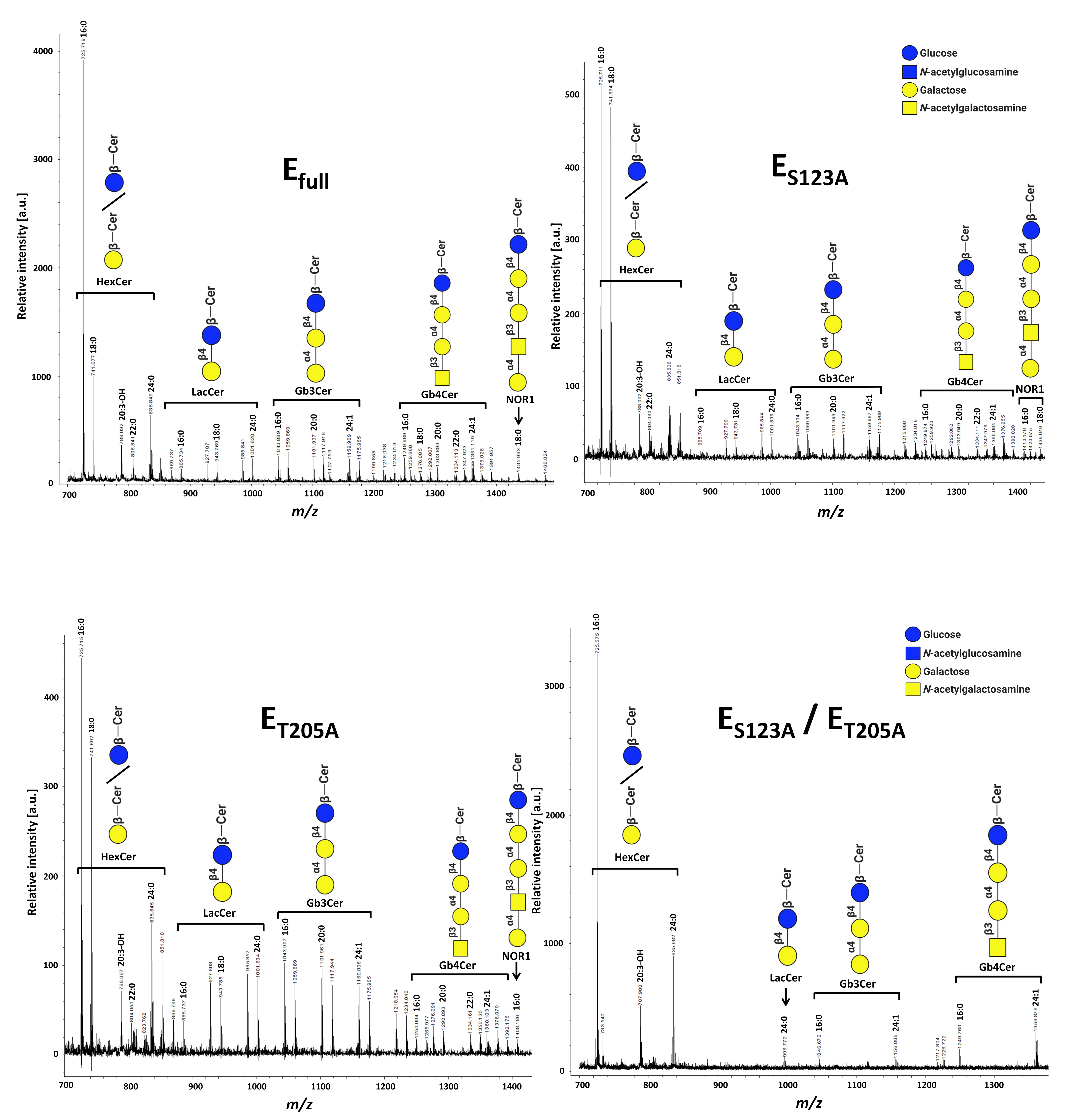

Fig. S3. **Reflectron-positive mode MALDI-TOF mass spectra of glycosphingolipids isolated from CHO-Lec2 cells expressing glycovariants of mutein.** The GSLs samples from CHO-Lec2 cells transfected with vectors encoding fully N-glycosylated E_full_ enzyme as well as E_S123A_, E_T205A_ and E_S123A_/E_T205A_ glycovariants.

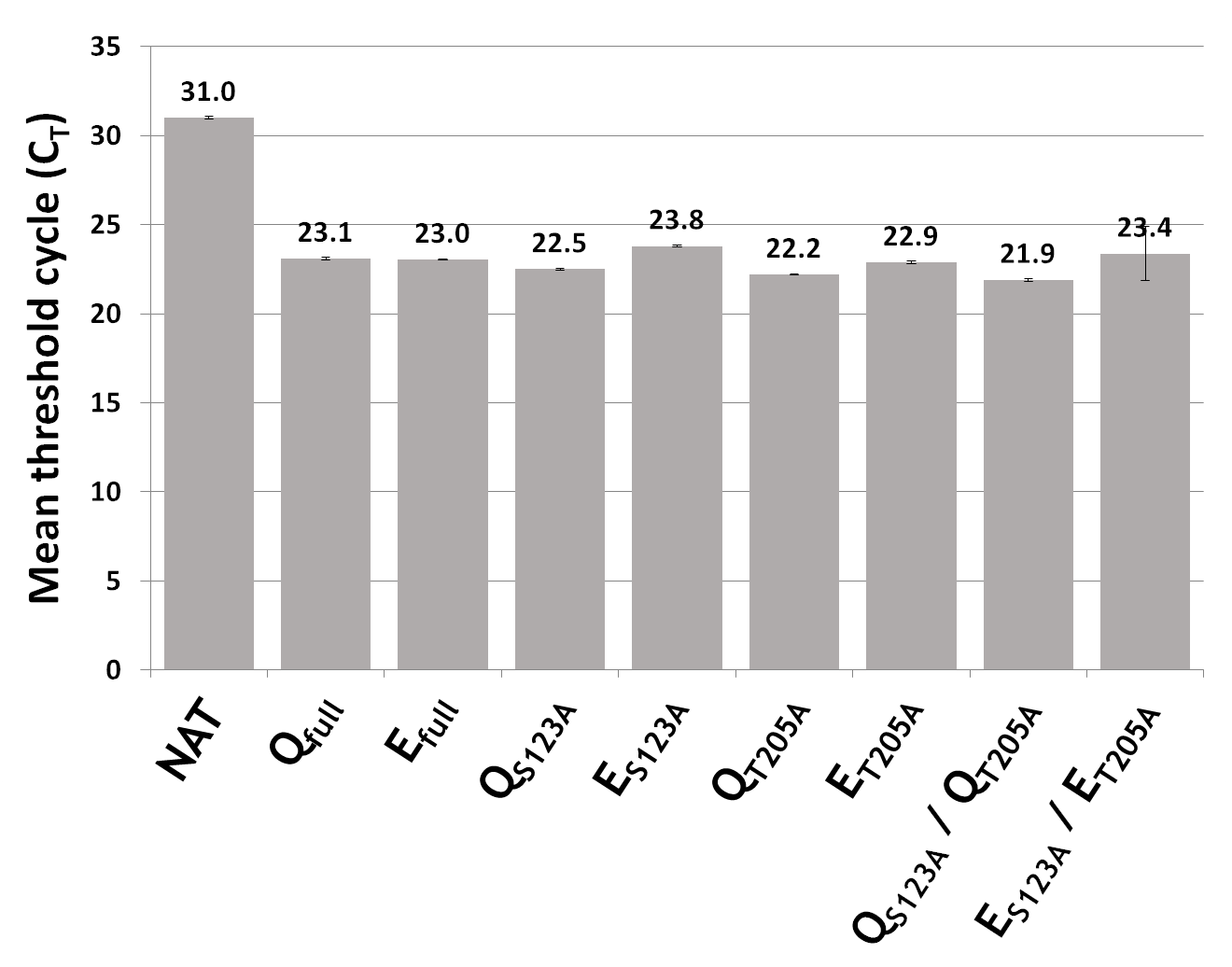

Fig. S4. **Comparisons of mean threshold cycle (C_T_)** **values between Gb3/CD77 synthase and its mutein glycovariants.** Q_full_, fully N-glycosylated of Gb3/CD77 synthase; E_full_, fully N-glycosylated mutein Gb3/CD77 synthase; Q_S123A_, Gb3/CD77 synthase with p.S123A substitution; E_S123A_, mutein Gb3/CD77 synthase with p.S123A substitution; Q_T205A_, Gb3/CD77 synthase with p.T205A substitution; E_T205A_, mutein Gb3/CD77 synthase with p.T205A substitution; Q_S123A_/Q_T205A_, Gb3/CD77 synthase with p.S123A/p.T205A substitutions; E_S123A_/E_T205A_, mutein Gb3/CD77 synthase with p.S123A/p.T205A substitutions.

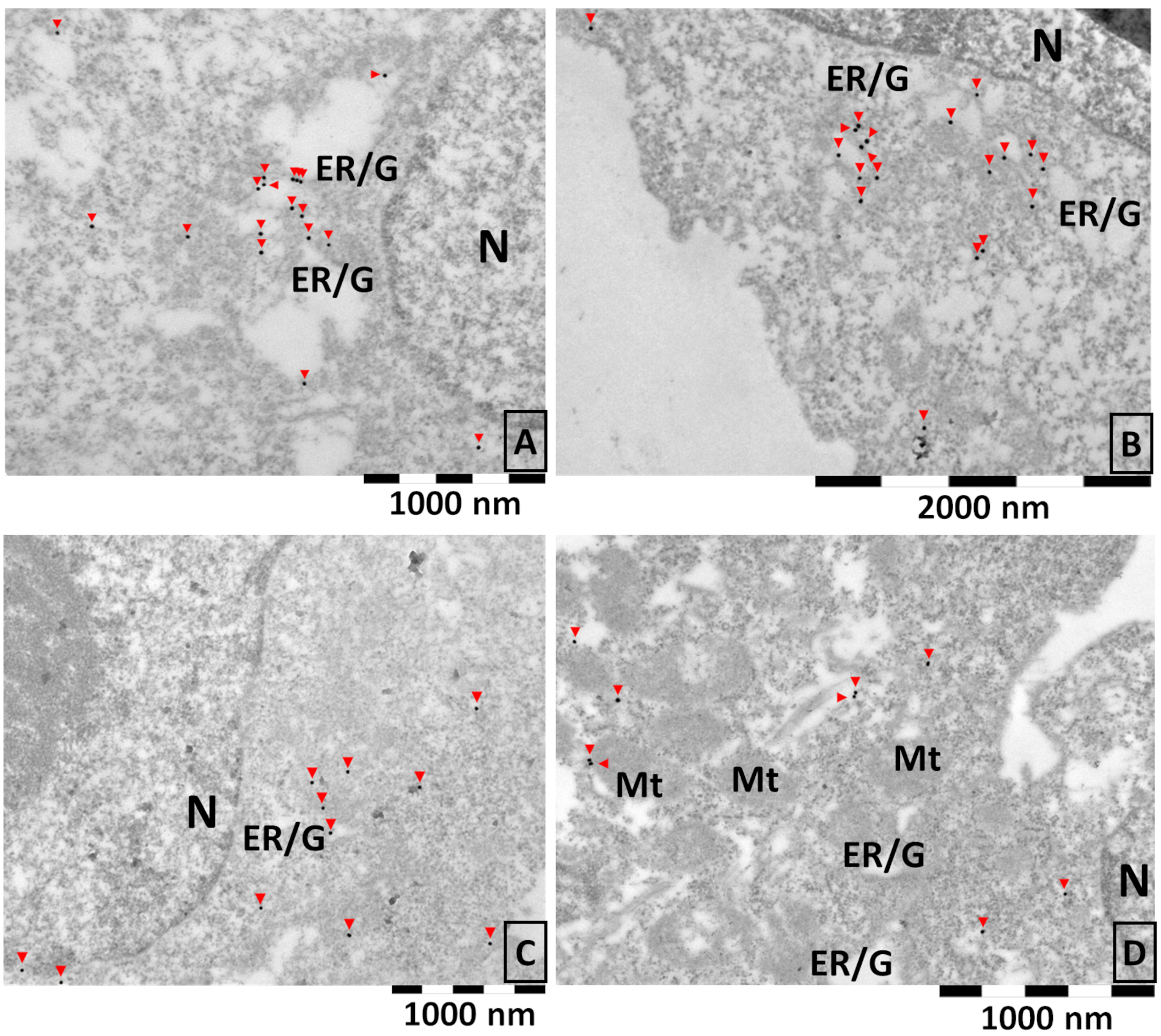

Fig. S5. **Subcellular localization of** **human Gb3/CD77 synthase glycovariants in CHO-Lec2 cells using immunogold reaction**. (**A**) Fully N-glycosylated Gb3/CD77 synthase. (**B**) Glycovariant Q_S123A_ with p.S123A substitution. (**C**) Glycovariant Q_T205A_ with p.T205A substitution. (**D**) Glycovariant Q_S123A_/Q_T205A_ with p.S123A/p.T205A substitutions. Red arrows indicated gold nanoparticles which correpond Gb3/CD77 synthase localization. N, nucleus; ER/G, endoplasmic reticulum or Golgi apparatus; Mt, mitochondrion.

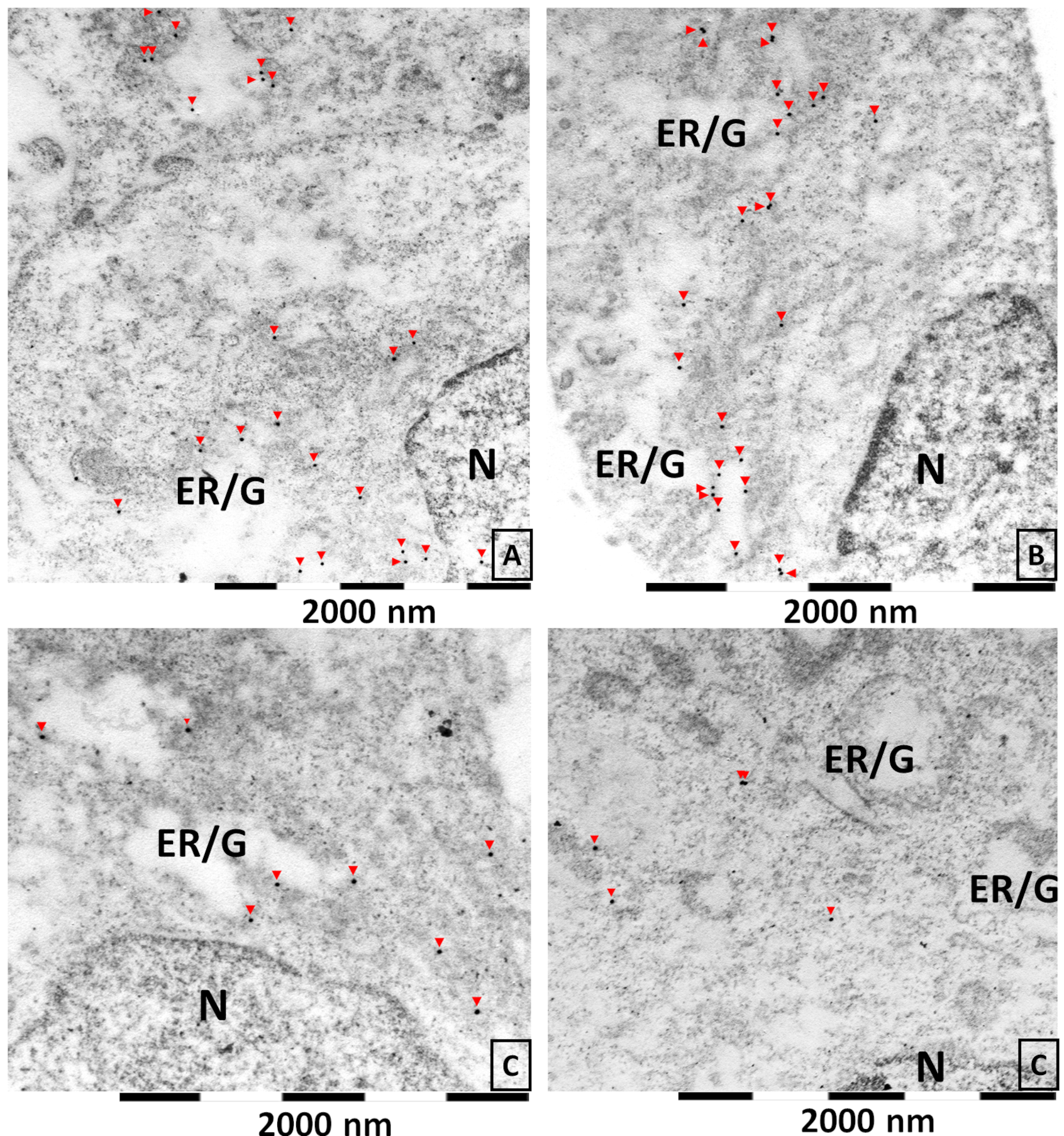

Fig. S6. **Subcellular localization of** **mutein** **glycovariants in CHO-Lec2 cells using immunogold reaction**. (**A**) Fully N-glycosylated Gb3/CD77 synthase. (**B**) Glycovariant E_S123A_ with p.S123A substitution. (**C**) Glycovariant E_T205A_ with p.T205A substitution. (**D**) Glycovariant E_S123A_/E_T205A_ with p.S123A/p.T205A substitutions. Red arrows indicated gold nanoparticles which correpond mutein localization. N, nucleus; ER/G, endoplasmic reticulum or Golgi apparatus.
